# Dysplastic Epithelial Repair Propagates Chronic Pathology Through the Paracrine Transformation of Pulmonary Fibroblasts

**DOI:** 10.64898/2026.04.02.716135

**Authors:** Nicolas P. Holcomb, Ronel Z. Samuel, Alena Klochkova, Joanna Wong, Sara Kass-Gergi, Meryl Mendoza, Michael M. Maiden, Evelyn A. Martinez, Diana M. Abraham, Madeline Singh, Harshini Kelam, Jarod A. Zepp, Andrew E. Vaughan

## Abstract

Severe lung injury promotes the ectopic accumulation of basal cells in the alveoli and the presence of these dysplastic epithelial cells are strongly associated with regions of pulmonary fibrosis (PF) in diseased lungs. Recent studies have identified a unique subset of “inflammatory” fibroblasts expressing pro-inflammatory genes, especially cytokines involved in monocyte recruitment, that are also enriched in disease and thought to contribute to the onset and progression of PF. Here we show that these two injury-induced cell types are intricately connected, in that dysplastic basal cells generate diffusible signals to robustly induce the inflammatory phenotype in pulmonary fibroblasts. Capitalizing on transcriptomic analysis, we identify the enriched inflammatory signaling pathways in treated fibroblasts and specifically demonstrate that IL-1α secreted by dysplastic basal cells is responsible for this fibroblastic transformation. IL-1α neutralization *in vivo* is sufficient to significantly reduce the inflammatory fibroblast burden in regions of alveolar bronchiolization, and the resolution of inflammatory fibroblasts in turn reduces CCR2+ immune cell recruitment to these areas. These results suggest dysplastic basal cells play an indirect role in chronic inflammation and fibrotic remodeling through the induction of a proinflammatory fibroblast phenotype and subsequent recruitment of immune cells, establishing a chronic wound healing microenvironment that prolongs localized pathologic remodeling.

## Introduction

Severe respiratory viral infections, including influenza and SARS-CoV-2, can destroy large regions of the alveolar epithelium and precipitate life-threatening complications including pneumonia and acute respiratory distress syndrome (ARDS) (*1–5*). Beyond acute lethality, there is increasing evidence that a significant portion of survivors of severe lung injury suffer long-term consequences, including pulmonary fibrosis (PF) (*6–8*). In most cases of lung injury, repair proceeds through a euplastic regenerative program primarily facilitated by the facultative stem cells of the alveoli, the alveolar type 2 cells (AT2s), restoring the alveolar niche and specialized architecture required for efficient gas-exchange (*9–12*). However, this regenerative response can be overwhelmed in cases of severe injury. Under these conditions, basal-like stem cells present in the airways proliferate and migrate distally to ectopically localize within the alveolar niche (*13–16*). While these ectopic KRT5+ basal cells act rapidly to prevent barrier failure and edema, they also occupy precious gas-exchanging surface area in the lung, persist chronically, and their strong bias towards an airway-fate results in alveolar bronchiolization, thus representing dysplastic repair (*17*, *18*). Critically, while inhabitation of alveolar space by dysplastic cells is itself detrimental to lung function, there is mounting evidence that these cells further propagate pathology through paracrine effects on other cell types within the distal lung (*19–21*).

Consistent with this possibility, expanded basal cell populations, including KRT5^neg^ p63-expressing “basaloid” cells in humans, are observed in fibrotic lung diseases as well as following severe viral injury (*8*, *18*, *20*, *22–25*). In non-resolving and fatal cases of SARS-CoV-2 infection, KRT5+ cells have been observed adjacent to fibrotic regions, suggesting a close spatial association between alveolar bronchiolization and fibrotic remodeling (*22*, *26*). Similarly, airway basal cell gene signatures have been found to correlate with mortality in cases of idiopathic pulmonary fibrosis (IPF), and increased quantities of basal cells have been associated with reduced forced vital capacity (*27*, *28*). Together, these observations raise the prospect that rather than just prognosticators of severe injury, ectopic KRT5+ basal cells are themselves drivers of chronic lung dysfunction. However, the functional consequences of basal cell expansion within the alveolar niche, and the mechanisms by which these cells influence surrounding cells, remain incompletely defined. We thus investigated the possibility that dysplastic bronchiolization reshapes the alveolar landscape through the introduction of a new signaling milieu into a region not adapted to interpret airway-derived cues.

Here we used an *in vivo* model of severe influenza-induced lung injury to initiate dysplastic repair and investigate regions of alveolar bronchiolization using spatial transcriptomics and immunofluorescent imaging, complemented by primary cell culture assays. We find that ectopic basal cells drive an inflammatory gene expression program in pulmonary fibroblasts through diffusible factors and identify IL-1α as the key basal cell-derived ligand mediating this response. This response is characterized by the upregulation of a distinct transcriptomic signature including expression of a variety of cytokines, chemokines, and immune related processes (*29–32*). In dysplastic regions, inhibition of the inflammatory fibroblast phenotype via IL-1α blockade was accompanied by an apparent decrease in monocyte and related CCR2+ immune cell occupancy. Together, this study highlights how dysplastic basal cells actively influence their ectopic niche through paracrine signaling to surrounding fibroblasts, inducing an inflammatory phenotype, and ultimately contribute to chronic inflammation and potentially PF.

## Results

### Dysplastic Regions are Enriched for Collagen Deposition and SAA3+ Fibroblasts

While we and others have demonstrated the long-term inhabitation of the alveolar parenchyma by basal-like cells after influenza injury, the functional consequences of their expansion within the alveolar niche and the mechanisms by which these cells shape the surrounding microenvironment have not been fully defined (*18*, *33–35*). To investigate this further, we infected Krt5-CreERT2 Ai14tdTomato transgenic mice with influenza (H1N1/PR8) and initiated lineage tracing on day 11 to ensure proper sampling of dysplastic regions (*36*, *37*). At 21 days-post-infection (DPI), we identified tissue sections bearing KRT5+ cells tdTomato expression), with prominent fibrotic regions readily identifiable in the hematoxylin and eosin (H&E) staining (Supplemental Figure 1). We then proceeded with spatial transcriptomics (Figure 1A). We applied both graph-based and K-means clustering to the spatial transcriptomics data and identified differentially expressed genes across clusters (Figure 1A; Supplemental Figure 1). We initially applied broad clustering resulting in four groups, and observed that clusters 2 and 3, marked by *Krt5* expression, also exhibited the highest levels of both *Col1a1* and *Saa3* (Figure 1A, 1B), features that were absent in healthy control lungs (Supplementary Figure 1). We confirmed COL1A1 protein deposition via immunofluorescence imaging, whereas regions populated by RAGE+ alveolar type 1 cells (AT1s) bore much less collagen (Figure 1C). While we hypothesized that regions of alveolar bronchiolization would be enriched for fibrotic markers, consistent with *Col1a1* expression, we also observed that *Saa3* was markedly enriched and largely confined to these regions of *Krt5* expression. (Figure 1B). SAA3 is a recently identified biomarker of an inflammatory fibroblast population hypothesized to orchestrate lung inflammation and fibrosis in a variety of lung diseases, though the exact function of these cells and whether they are primed to become fibrotic or drive immune cell recruitment is unclear (*29–32*). We confirmed the long-term association between dysplastic KRT5+ epithelia and SAA3+ fibroblasts through at least 50 DPI, suggesting potential signaling interactions between these populations. (Figure 1D). To assess whether alveolar fibroblasts preferentially adopt the inflammatory fibroblast state within regions of alveolar bronchiolization, we performed lineage tracing with Scube2-CreERT2 mice which specifically labels alveolar fibroblasts (*31*). Tamoxifen was administered to initiate lineage-tracing two weeks prior to infection with influenza and lungs were harvested 21 DPI followed by quantification of the total number of SAA3+ inflammatory fibroblasts (PDGFRα+) within KRT5+ regions that bore the lineage trace. Within dysplastic regions, we identified populations of both Scube2-lin derived and lineage-negative PDGFRα+ inflammatory fibroblasts (Figure 1E). Furthermore, we observed no significant reduction in Scube2-lin fibroblasts within dysplastic regions 21 DPI compared to uninfected controls (Figure 1F). Notably, Scube2-lin derived cells accounted for an average of 45% of the inflammatory fibroblasts in the KRT5+ regions (Figure 1G). Given that this tracing scheme accounts for ∼57% of PDGFRα+ fibroblasts in uninjured lungs, this suggests that additional fibroblast lineages (adventitial, peribronchial) may also contribute to the pool of inflammatory fibroblasts in these areas. Overall, we observed a proximity-dependent enrichment of the lineage-traced inflammatory fibroblast population within dysplastic regions (Supplemental Figure 2). This spatial distribution suggests a localized mesenchymal response, in which fibroblasts closer to the dysplastic KRT5+ epithelia are primed into an inflammatory state and hence drive immune recruitment.

**Figure 1:**
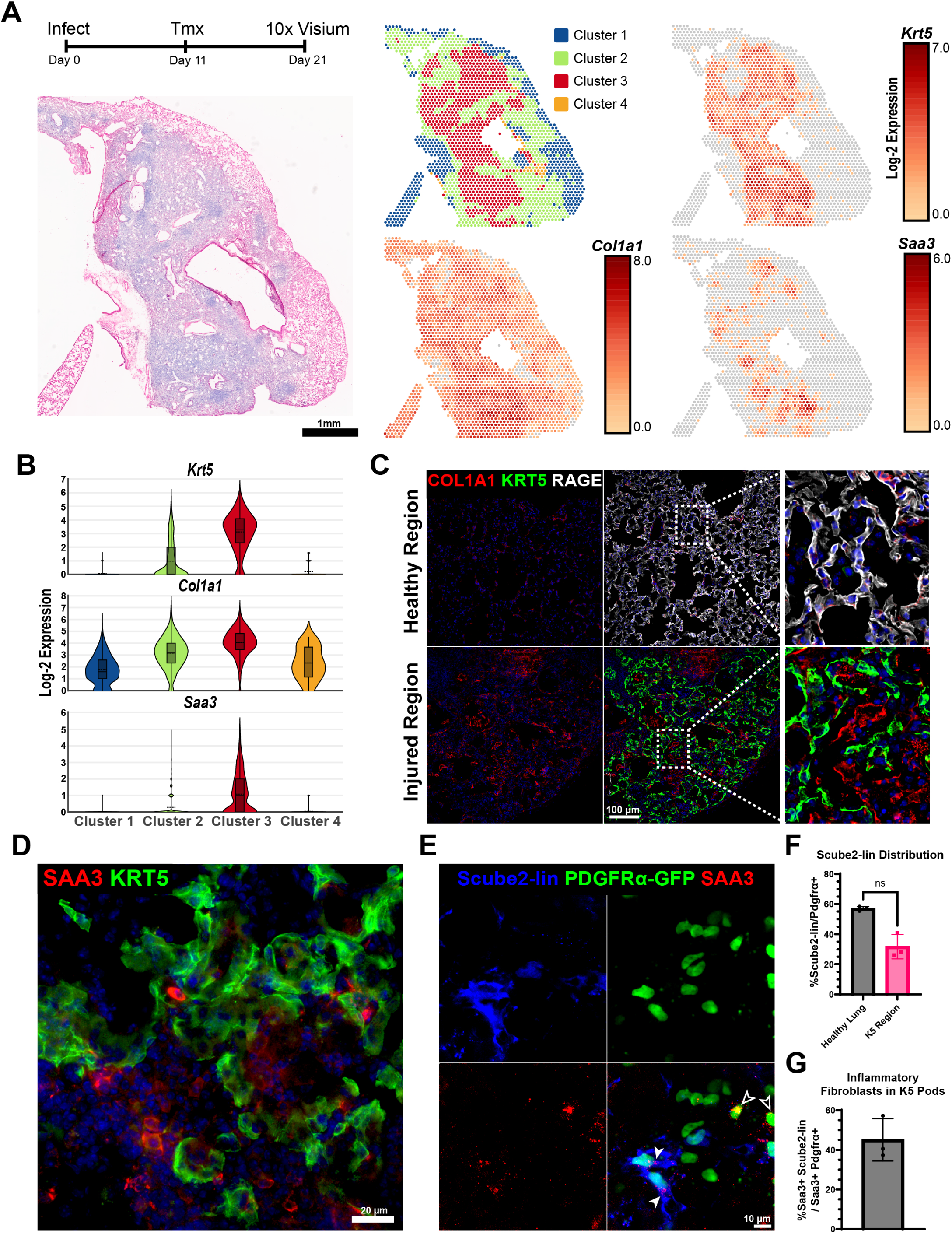
Regions of Alveolar Bronchiolization are Enriched for COL1A1 Content and SAA3+ Fibroblasts. (**A**) Schematic showing the timing of infection, induction of lineage trace (Krt5-CreERT2), and tissue harvest for spatial transcriptomics alongside an H&E Image of the full 21-days post-infection (DPI) selected lobe. Spatial sequencing plots of the clustering and the log-2 expression for *Krt5*, *Col1a1*, and *Saa3* are on the right. (**B**) Log-2 expression of the indicated genes within each cluster in (A) as violin plots. (**C**) Representative immunofluorescence images from a 21-DPI mouse showing a healthy region populated by alveolar type 1 cells (RAGE+) and a region of alveolar bronchiolization, populated by ectopic basal cells (KRT5+). Insets highlight the difference in COL1A1 content. 20x images, DAPI (blue), COL1A1 (red), KRT5 (green), RAGE (white). (**D**) Representative image from a 50-DPI mouse highlighting SAA3+ cells in close proximity to regions populated by ectopic basal cells (KRT5+), DAPI (blue), SAA3 (red), KRT5 (green). **(E)** Representative immunofluorescence image from a Scube2CreERT2; PDGFRα-H2B:EGFP; Ai14tdTomato 21-DPI mouse showing Scube2-lineage-traced SAA3+ fibroblasts within a region populated by ectopic basal cells (KRT5+), Scube2-lineage (blue), PDGFRα (green), SAA3 (red). **(F)** Quantification of the Scube2-lin fibroblasts as a percentage of total PDGFRα+ cells in the lungs of healthy mice and within KRT5+ dysplastic regions at 21 DPI. **(G)** Proportion of inflammatory fibroblasts within KRT5+ regions derived from Scube2-lin alveolar fibroblasts. (B) Data shows log-2 expression within the 4 clusters in (A) as volcano plots; min, q1, mean, median, q3, max. (E) Closed arrow heads: Scube2-lin inflammatory fibroblasts. Open arrow heads: lineage negative PDGFRα+ inflammatory fibroblasts. (F) Percentage of Scube2-traced cells out of total PDGFRα+ fibroblasts ± SD (n=3 mice per group). (G) Percentage of SAA3+ Scube2-traced cells out of total SAA3+ PDGFRα+ fibroblasts ± SD (n=3 mice). *P* values were calculated using two tailed Mann-Whitney U test: ns: not significant.

### Ectopic Krt5+ Cells Induce a Direct Inflammatory Phenotype on Pulmonary Fibroblasts

Next, we assessed whether factors secreted by ectopic KRT5+ basal cells are sufficient to directly drive *Saa3* expression in pulmonary fibroblasts. To this end, we generated conditioned media (hereafter CM) from post-influenza ectopic KRT5+ basal cells expanded *in vitro* (Figure 2A). Recognizing the potential impact of matrix stiffness of fibroblast behavior, CM or control media was used to treat pulmonary fibroblasts on dishes of varying stiffness: plastic, 16 kPa, and 0.5 kPa. The 16 kPa dishes reflect the native stiffness of the bronchi while 0.5kPa dishes more accurately reflect the stiffness of the alveoli (*38*). CM treatment induced a significant upregulation of key inflammatory fibroblast marker genes *Saa3* and *Ccl2* across plating conditions and the cells appeared increasingly sensitive to stimulation on softer, more physiologically relevant, dishes (Figure 2B). To gain further insight into this inflammatory fibroblast phenotype, we performed bulk RNA-seq on control and CM-stimulated fibroblasts cultured on 0.5 kPa or plastic dishes (Figure 2C, Supplemental Figure 3). We observed a large number of significant (p<0.01) and highly upregulated (>4 fold) cytokines and chemokines upon CM treatment (Figure 2D). We noted dramatic and significant upregulation of inflammatory (*Saa3, Lcn2, Cxcl5, IL6, Ccl2*) and proliferative (*Mki67, Top2a*) marker genes alongside a modest downregulation of fibrotic (*Cthrc1, Col1a1, Postn*) marker genes (Figure 2E). We performed a functional enrichment analysis of the top 92 significantly upregulated genes shared between 0.5kPa and Plastic CM treated fibroblasts using Metascape (*39*), which highlighted enrichment of many immune-related processes, inflammatory signaling pathways, and an association with lung fibrosis (Figure 2F). The second most significant enrichment category was for the cellular response to interleukin-1 (IL-1), and notably recombinant IL-1β is known to be sufficient to induce the inflammatory fibroblast phenotype in culture (*31*, *40*).

**Figure 2:**
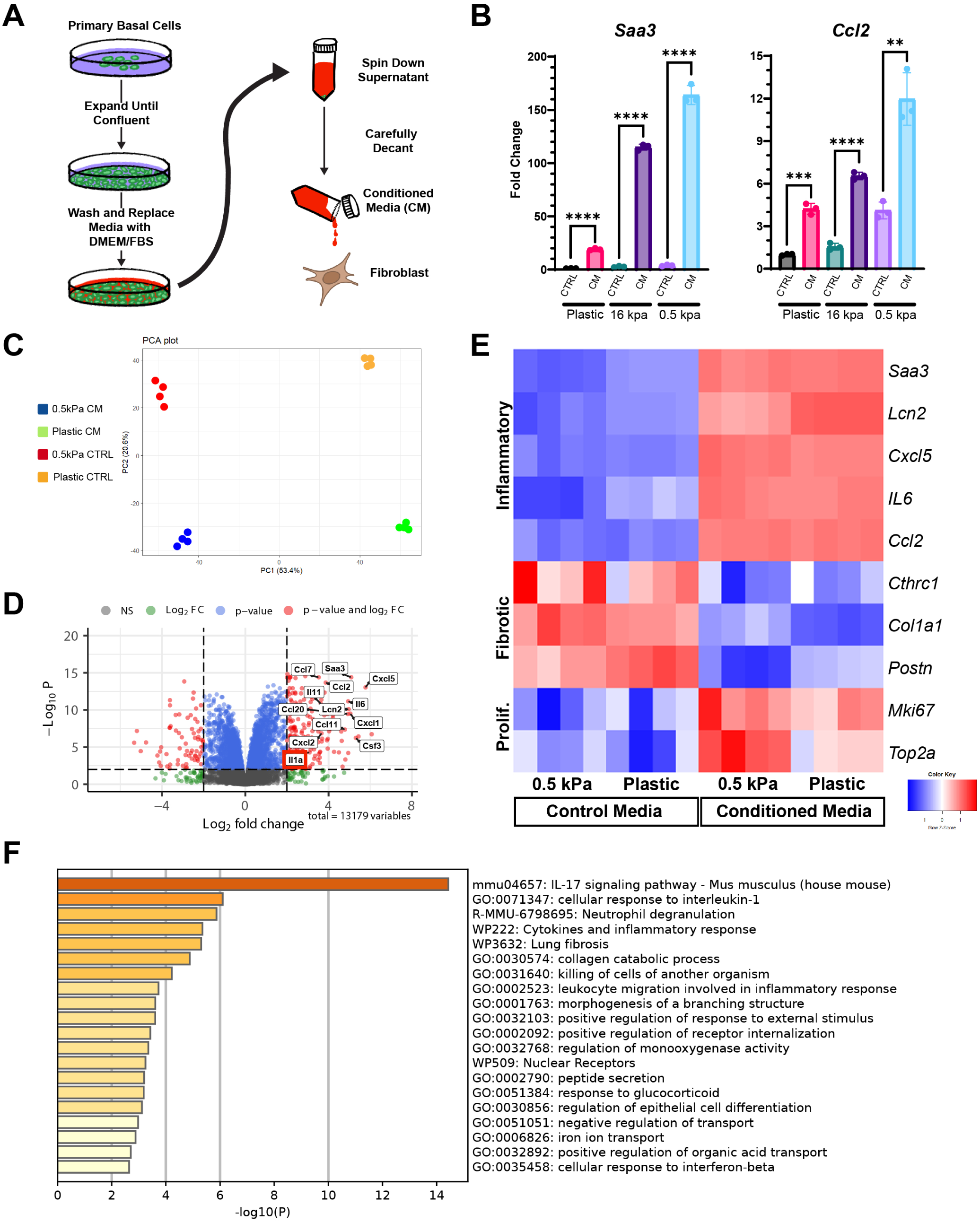
Dysplastic Basal Cells Induce an Inflammatory Phenotype in Pulmonary Fibroblasts *in vitro* via Diffusible Factors. (**A**) Graphic summarizing the generation of conditioned media from ectopic, post-influenza KRT5+ basal cells used to treat fibroblasts. (**B**) Key inflammatory fibroblast marker genes *Saa3* and *Ccl2* expression was assayed by RT-qPCR for control and conditioned media treated fibroblasts on plastic, 16kPa, and 0.5kPa plating conditions. Plotted as fold change relative to Plastic Control. (**C**) PCA Plot showing the 16 samples used for bulk RNA-seq analysis. (**D**) Volcano plot illustrating significantly differentially expressed genes between control and conditioned media treated fibroblasts cultured on 0.5kPa plates with selected cytokines and chemokines labeled. *P* value < 0.01, Fold Change > 4,184 upregulated 68 downregulated. We note upregulation of *Il1a* in the fibroblasts themselves upon conditioned media treatment. (**E**) Heatmap depicting the relative expression of selected inflammatory, fibrotic, and proliferative fibroblast marker genes between the samples depicted in (C). (**F**) Functional enrichment analysis of the top 92 significantly upregulated genes (*P* value < 0.01, Fold Change > 4) shared between 0.5kPa and Plastic conditioned media treated fibroblasts via Metascape. (B) Data shows mean fold change ± SD and are pooled from 3 independent experiments (n=3 mice). *P* values were calculated using unpaired *t*-tests: \*\**P* < 0.01, \*\*\**P* < 0.001, \*\*\*\**P* < 0.0001.

To further identify diffusible factors secreted by KRT5+ ectopic basal cells into the CM, we employed a membrane-based antibody array to determine relative levels of selected cytokines and chemokines. The ratio of protein in the CM relative to the control media was quantified and we identified a number of interesting candidates at >10x higher concentrations in the CM (Supplemental Figure 4). However, fibroblasts treated with many of these candidates including GDF15, VEGF, OPN, CXCL1, IGFBP3, TNFRSF11B, LIX, TF, or CCL20 failed to induce the inflammatory gene expression profile demarcated by *Saa3* (Supplemental Figure 4). Interestingly, despite pathway identification from our RNA-Seq experiments, neither IL-1α nor IL-1β were present at high enough concentrations in CM to be detected using this antibody array.

### IL-1α Drives Inflammatory Fibroblast Activation

Despite our inability to detect either IL-1 cytokine in the array format, IL-1 is known to possess activity at extremely low levels. To discern whether low level IL-1 might still be responsible for the observed phenotype, we devised a gamut of *in vitro* experiments leveraging receptor antagonists, recombinant proteins, and neutralizing antibodies. First, we treated fibroblasts with CM supplemented with varying concentrations of IL-1 receptor antagonist (IL1RA), which binds to the IL-1 receptor (IL1R) blocking both IL-1α and IL-1β binding and signal transduction. Strikingly, we observed that IL1RA supplementation was sufficient to block the induction of the inflammatory phenotype upon CM treatment, indicating that the IL1R is necessary for this process and therefore confirming that the cellular response to IL-1 is driving this phenotype (Figure 3A). IL-1β has been shown to induce the inflammatory fibroblast phenotype in prior publications, but IL-1β is produced largely by myeloid cells not present in our *in vitro* cultures (*31*, *40–43*). Accordingly, we reasoned that IL-1α was more likely to be present in CM (Supplemental Figure 5). To interrogate whether IL-1α was the ligand responsible, we treated fibroblasts with control media, control media supplemented with recombinant IL-1α (rIL-1α), CM, and CM supplemented with IL-1α neutralizing antibodies (nAb). rIL-1α supplemented in the control media alone was sufficient to strongly drive the inflammatory fibroblast phenotype, while IL-1α nAb supplemented in the CM was sufficient to completely block *Saa3* induction, indicating that IL-1α is the ligand responsible for driving this phenotype (Figure 3B). However, we also noted significant upregulation of IL-1α in the CM treated fibroblasts themselves suggests that the fibroblasts themselves may be the actual source of IL-1α, produced downstream of some other factor in the CM (Figure 2D – Red box). Thus, while IL-1α was strictly necessary, it could be driving this phenotype in either a paracrine manner through secretion by the ectopic basal cells themselves or could be produced in an autocrine fashion by fibroblasts downstream of some factor “k” secreted by ectopic basal cells (Figure 3C). We proceeded to investigate these possibilities.

**Figure 3:**
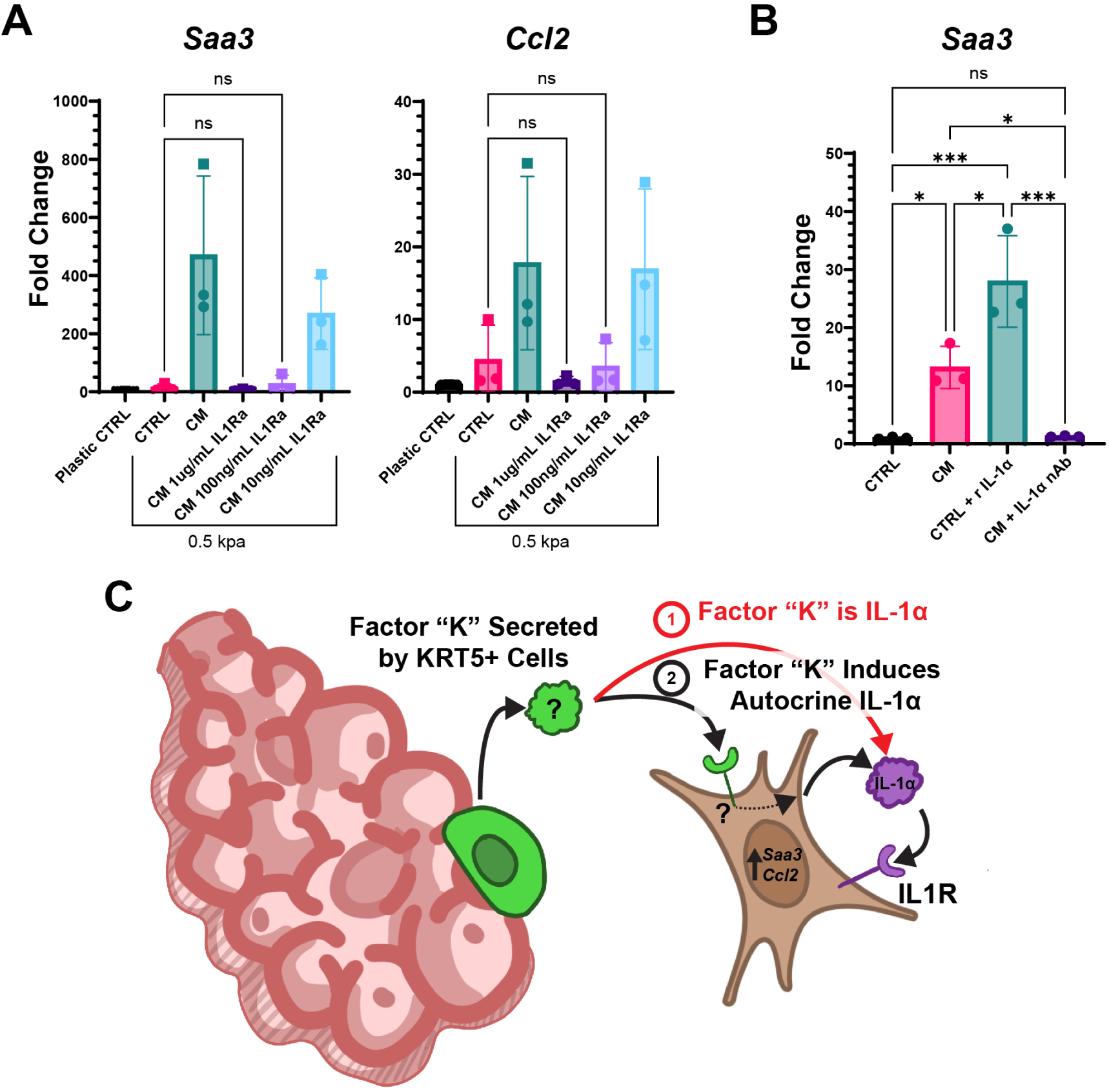
IL-1α Drives Inflammatory Fibroblast Phenotype Following Conditioned Media Treatment. (**A**) IL-1 receptor blockade with recombinant IL1RA completely prevents *Saa3* and *Ccl2* upregulation in fibroblasts induced by conditioned media, as assessed by RT-qPCR for fibroblasts cultured on 0.5kPa plates. Media was supplemented with recombinant IL1RA at 1 µg, 100ng, and 10ng/mL concentrations with fold change relative to plastic control. (**B**) rIL-1α at 50ng/mL added to control media or IL-1α neutralizing antibodies at 100ng/mL added to conditioned media with fold change relative to control. (**C**) Graphic depicting the potential sources of IL-1α driving the inflammatory phenotype. [(A) and (B)] Data shows mean fold change ± SD and are pooled from 3 independent experiments (n=3 mice). *P* values were calculated using either [(A) and (B)] a one-way ANOVA with post-hoc Turkey multiple comparison test. \**P* < 0.05, \*\**P* < 0.01, \*\*\**P* < 0.001, ns: not significant.

### Ectopic KRT5+ Basal Cell Derived IL-1α Drives the Inflammatory Fibroblast Phenotype

To determine the cellular source(s) of IL-1α driving the inflammatory fibroblast phenotype, we generated CM from our control and CM-stimulated fibroblasts themselves as well as ectopic basal cells, then measured and quantified the concentration of IL-1α present through an ELISA. Despite induction at the RNA level (Fig. 2D), stimulated fibroblast IL-1α levels were below the limit of detection. Conversely, the concentration of IL-1α in ectopic basal cell CM was quantified at 2.63 pg/mL (Figure 4A). While IL-1α is known to be active within the picogram range, this concentration was still very low and necessitated additional confirmation that it is sufficient to drive the inflammatory fibroblast phenotype (*43*). To this end, we tested dosing around the aforementioned concentration of IL-1α, demonstrating that the quantified level of rIL-1α was sufficient to induce this phenotype (Figure 4B). To confirm that IL-1α from the ectopic basal cells is driving the inflammatory phenotype and investigate whether other secreted ligands contribute, we performed a series of antibody depletion experiments. We incubated control media or CM with IL-1α nAb followed by immunodepletion via column-based magnetic separation (Figure 4C). This approach resulted in a strong reduction of the inflammatory fibroblast phenotype which was not further reduced via “spiking in” of additional IL-1α nAb during fibroblast treatment. We therefore conclude that ectopic basal cell-derived IL-1α is primarily responsible for driving the inflammatory fibroblast phenotype (Figure 4D). Interestingly, we did observe small but significant remaining upregulation of *Saa3 and Ccl2* relative to control with CM even after IL-1α depletion, suggesting that there may be additional (but much less active) secreted factors that contribute to driving the inflammatory phenotype.

**Figure 4:**
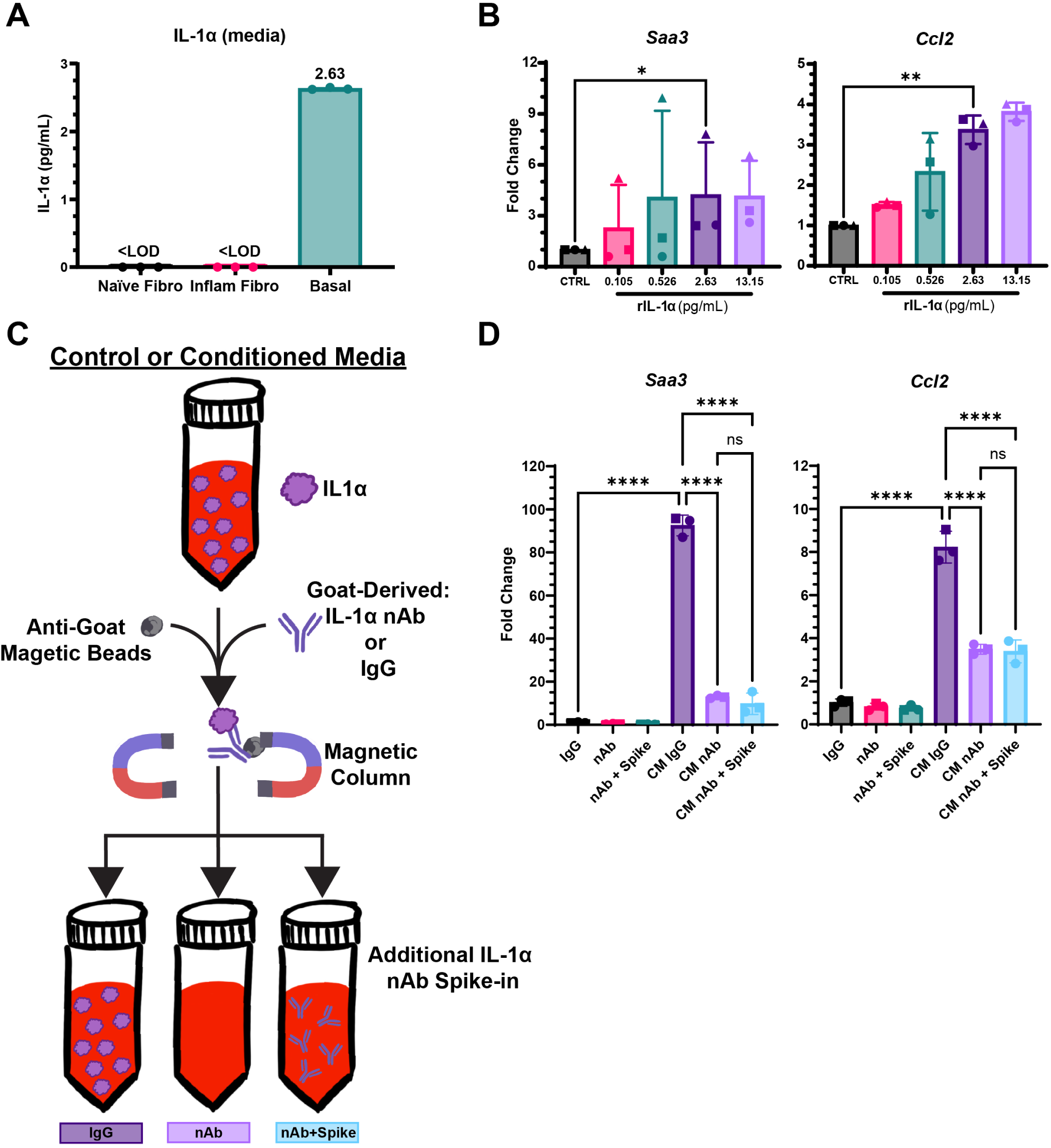
Basal Cell Derived-IL-1α Is Sufficient To Drive the Inflammatory Fibroblast Phenotype. (**A**) Quantified levels of IL-1α from naïve fibroblast (Naïve fibro), inflammatory fibroblast (Inflam fibro), and ectopic basal cell (Basal) conditioned media as determined by ELISA. (**B**) *Saa3* and *Ccl2* gene expression by RT-qPCR for fibroblasts cultured on 0.5kPa plates and treated with a 5-fold titration of IL-1α added to control media, including the level determined in (A). (**C**) Graphic illustrating the strategy used to deplete conditioned media of endogenous IL-1α with antibodies, magnetic beads, and column-based separation. (**D**) *Saa3* and *Ccl2* gene expression by RT-qPCR for fibroblasts cultured on 0.5kPa plates in control or conditioned media depleted using IgG (CTRL IgG or CM IgG) or IL-1α nAb (CTRL nAB, CM nAb) and supplemented with additional IL-1α nAb after depletion (+Spike). [(B) and (D)] Data shows mean fold change ± SD and are pooled from 3 independent experiments (n=3 mice). *P* values were calculated using either (B) a ratio paired *t*-test or (D) a one-way ANOVA with post-hoc Turkey multiple comparison test: \**P* < 0.05, \*\**P* < 0.01, \*\*\*\**P* < 0.0001, ns: not significant.

Inflammatory fibroblasts have been proposed to represent an intermediate step toward fully fibrotic fibroblasts, though this has only been inferred from pseudotime analyses (*31*). We therefore explored whether induction of the inflammatory fibroblast phenotype through rIL-1α was sufficient to sensitize these fibroblasts to subsequent fibrotic queues via titration of TGF-β. We found that 24-hour pre-treatment with rIL-1α did not significantly alter *Col1a1* or *Cthrc1* expression in response to TGF-β treatment at 1, 2, or 4 ng/mL (Supplemental Figure 6), arguing against inflammatory fibroblasts serving as a required intermediate toward fibrotic transformation, though leaving open the possibility that inflammatory fibroblasts contribute to fibrosis indirectly by modulating inflammation.

### IL-1α Blockade post-Influenza Reduces Inflammatory Fibroblast Burden in Dysplastic Regions

Because IL-1α serves as the primary driver of the inflammatory phenotype *in vitro*, we asked whether IL-1α blockade *in vivo* would be sufficient to inhibit fibroblasts from assuming an inflammatory phenotype within regions of alveolar bronchiolization. To address this question, we infected a cohort of mice with influenza, and delivered IL-1α nAb in PBS through daily Intraperitoneal (IP) injections from 45-49 DPI followed by tissue harvest at 50 DPI for downstream analysis (Figure 5A). The area of SAA3 within KRT5+ regions was analyzed through immunofluorescence, subsequently the relative area of SAA3 (Saa3 Area/KRT5 Area) was calculated. As predicted by *in vitro* modeling, we found that this short treatment with IL-1α nAb was sufficient to significantly reduce the SAA3 enrichment in these regions of bronchiolization (Figure 5B, 5C).

**Figure 5:**
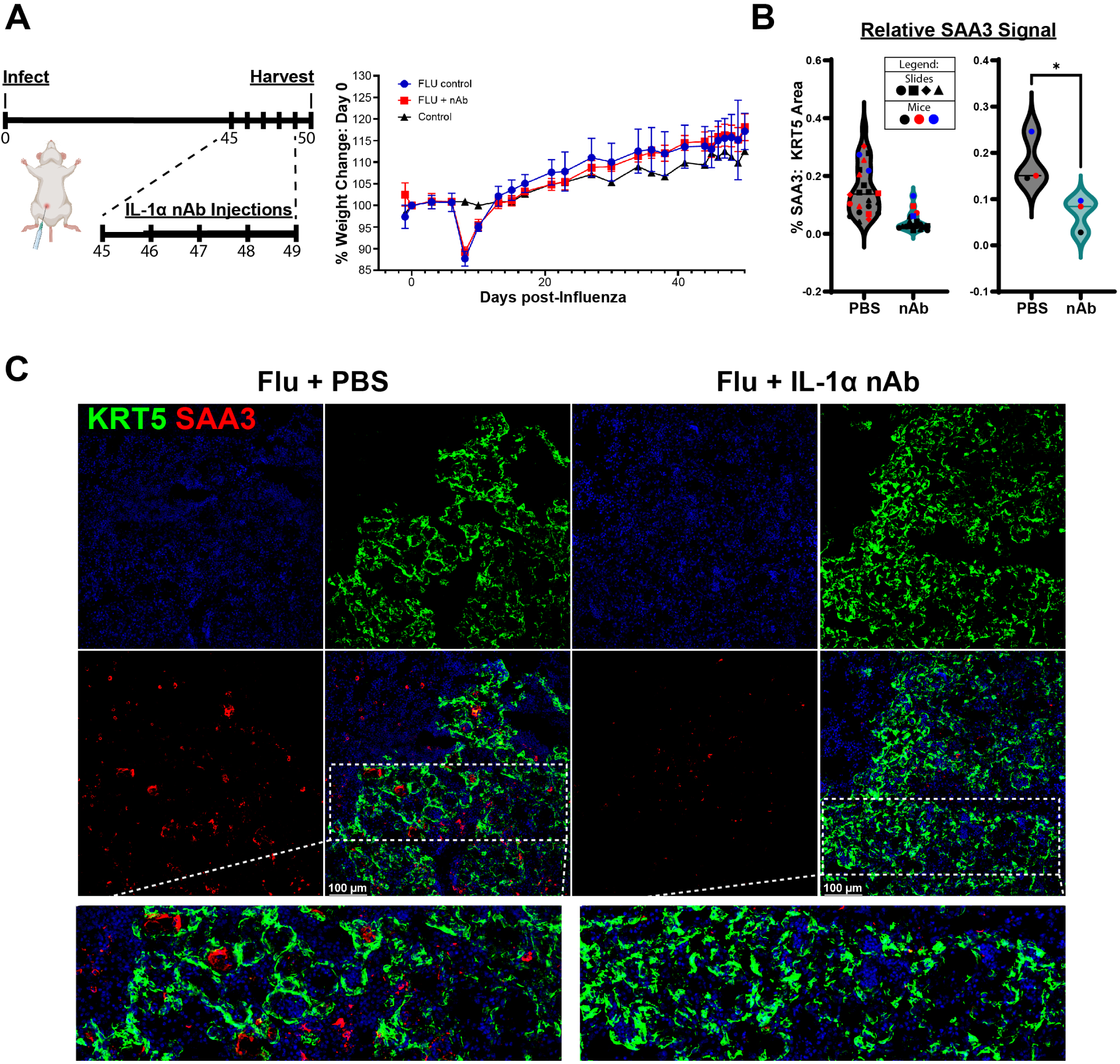
IL-1α Neutralization *in vivo* Significantly Reduces SAA3 Enrichment in KRT5+ Regions Post-Influenza. (**A**) Diagram showing the timing of the flu infection experiment and five IL-1α nAb injections (200 µg per injection). Average weight change for the three groups over the course of the course of 50 days is displayed on the right. (**B**) Violin plots showing the relative amount of SAA3 signal in KRT5+ Regions for flu infected mice that received PBS or IL-1α nAb. The graph on the left displays the quantified individual images and the graph on the right displays the averages per mouse (used for statistical analysis). (**C**) Representative immunofluorescence images from flu infected mice that received PBS or IL-1α nAb with insets highlighting the difference in SAA3 signal, DAPI (Blue) KRT5 (Green) SAA3 (Red). (B) Data shows average relative SAA3 signal (n=3 mice per group). A single outlier was identified using a two-sided Grubbs’ test and removed. *P* values were calculated using an unpaired *t*-test: \**P* < 0.05.

Given the significant reduction of SAA3 staining within KRT5+ regions upon IL-1α nAb delivery, we were interested if the suppression of the inflammatory fibroblast phenotype might reduce immune cell recruitment in these regions. Specifically, we hypothesized that fibroblasts driven to an inflammatory phenotype by IL-1α may propagate the CCL2-mediated recruitment of immune cells (e.g. monocytes and some macrophages) to these regions of alveolar bronchiolization (Figure 6A). CCL2 is known to drive the recruitment of bone marrow derived monocytes to the lung during injury, and we observed KRT5+ regions with CCR2+ cells in close proximity (Figure 6B) (*44–46*). Animals treated as above with IL-1α nAb exhibited an clear, strong trend toward decreased numbers of CCR2+ cells relative to KRT5 area (#CCR2/KRT5 area) (Figure 6C, 6D). Taken together, these findings suggest the inflammatory fibroblast phenotype contributes to CCR2+ immune cell recruitment and the establishment of a “chronic wound healing” microenvironment within sites of alveolar bronchiolization.

**Figure 6:**
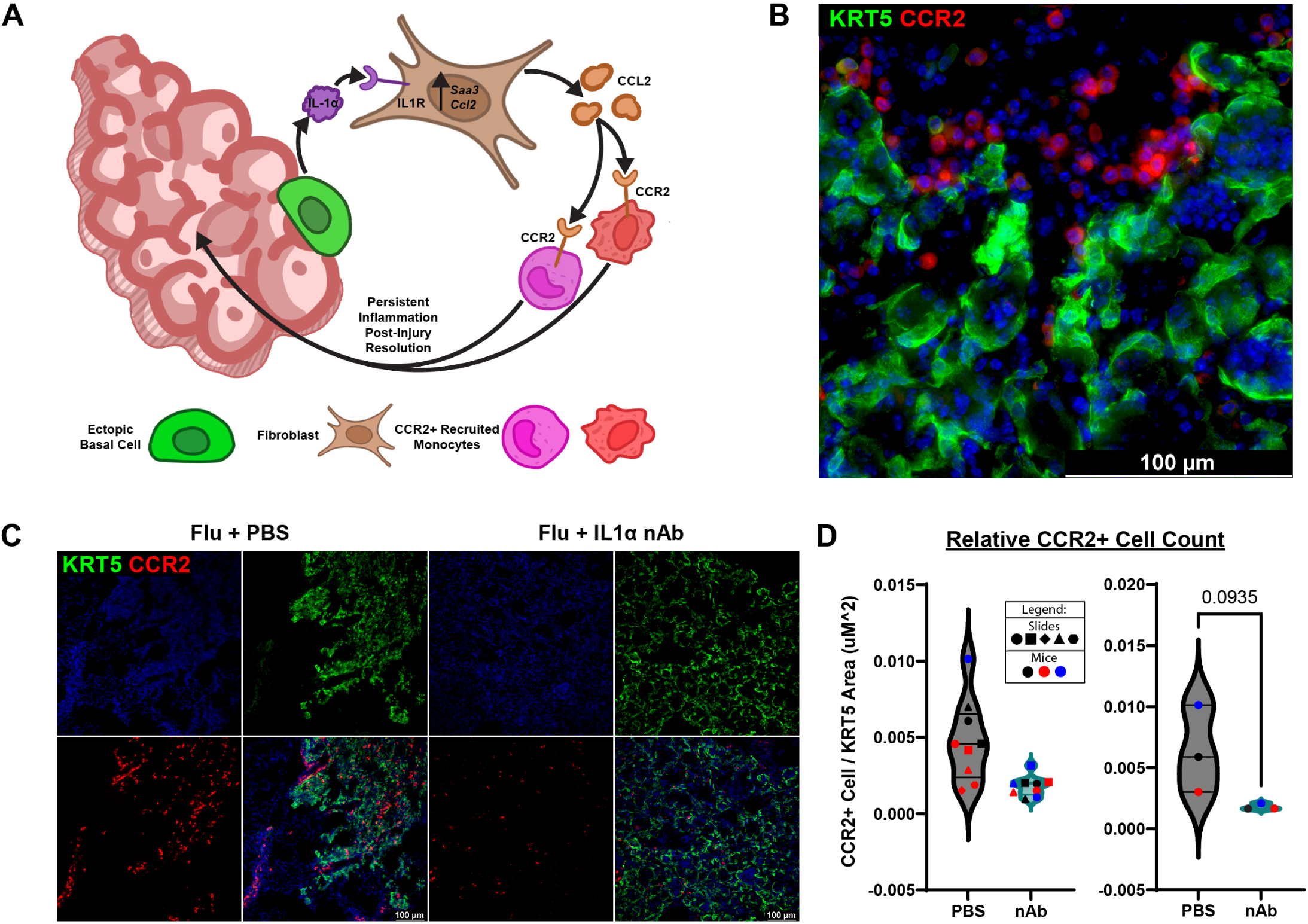
IL-1α Neutralization Correlates with Reduced CCR2+ Cell Enrichment in KRT5+ Regions. (**A**) Graphic showing a model where ectopic KRT5+ cells secrete IL-1α, driving surrounding fibroblasts towards an inflammatory phenotype, which results in the secretion of CCL2 by the fibroblasts, and persistent accumulation and retention of CCR2+ immune cells after resolution of injury. (**B**) Representative immunofluorescence image highlighting CCR2+ cell accumulation in close proximity to KRT5+ cells post-flu. (**C**) Representative immunofluorescence images from flu infected mice that received PBS or IL-1α nAb. 20x images, DAPI (Blue) KRT5 (Green) CCR2 (Red). (**D**) Violin plots showing the relative number of CCR2+ cells in KRT5+ Regions for flu infected mice that received PBS or IL-1α nAb. The graph on the left displays the quantified individual images and the graph on the right displays the averages per mouse (used for statistical analysis). (D) Data shows average relative CCR2+ Cell number (n=3 mice per group) as a violin plot. *P* values were calculated using an unpaired *t*-test.

## Discussion

While dysplastic repair presumably aids in rapid barrier restoration to limit pulmonary edema and protects against further insults acutely, the long-term maladaptive consequences result in reduced lung function and are associated with fibrotic lung diseases (*8*, *26*, *28*). Regions of dysplastic alveolar bronchiolization represent ectopic airway niches where KRT5+ basal cells proliferate, differentiate into a variety of airway cell types, and secrete atypical signals into their environment (*17*, *18*). These regions are largely non-resolving, at least in mice, where dysplastic patches have been identified one year post-infection (*18*, *35*). Here we highlight another layer of pathologic consequences resulting from paracrine / juxtacrine interactions between this persistent dysplastic epithelia and the surrounding tissue. This work provides insight into the spatial impact of dysplasia / bronchiolization and the cell-to-cell communication between ectopic KRT5+ basal cells and pulmonary fibroblasts. We specifically demonstrate that ectopic KRT5+ cells induce an IL-1α-dependent inflammatory phenotype in pulmonary fibroblasts. These inflammatory fibroblasts exhibit an upregulation of immune-related signaling processes which results in recruitment of CCR2+ immune cells. As such, this work highlights an important role for dysplastic epithelia in propagating tissue pathology via an epithelial-stromal-immune axis.

Our work adds a novel IL-1α dependent signaling axis to the growing evidence that basal cells signal to surrounding fibroblasts in their ectopic niches after viral injury and that there are chronic pathologic consequences to the dysplastic repair process (*20*, *21*, *47–51*). Many studies have provided valuable insight into the transcriptional profile of basal cells, but the broader cellular landscape within regions of alveolar bronchiolization has been less fully investigated (*18*, *24*, *52–56*). The advent of spatial transcriptomics presented an opportunity to overcome these limitations and assess how alveolar bronchiolization impacts tissue architecture and function. Similar to reports describing the association between ectopic basal / basaloid cells in fibrotic regions in patients with fibrotic lung disease and post-acute COVID-19 lung disease, our spatial transcriptomics data demonstrates an enrichment in fibrosis-associated signatures following severe influenza-induced lung injury in mice (Supplemental Figure 1). Although this association does not establish causality, it prompted us to investigate direct and indirect mechanisms by which ectopic KRT5+ basal cells may contribute to the propagation of PF and other deleterious changes to the tissue microenvironment.

Inflammatory fibroblasts have been identified in numerous pulmonary injury and disease states, though aside from IL-1β the discrete drivers of this inflammatory state have not been fully elucidated (*29–32*, *57*, *58*). Perhaps more importantly, the (patho)physiologic consequences of fibroblasts adopting said state are not clearly defined. Our studies provide additional evidence that a persistent inflammatory fibroblast burden may drive pathology through the sustained recruitment of immune cells. The significant reduction of inflammatory fibroblasts in dysplastic regions after IL-1α nAb treatment was accompanied by a strong trend towards decreased CCR2+ cell occupancy, supporting a direct role for inflammatory fibroblasts mediating immune cell recruitment. If chronic occupancy of the distal lung by ectopic KRT5+ cells and their progeny influence surrounding fibroblasts to recruit immune cells, it raises the possibility of a feed-forward circuit in which continuous immune cell recruitment leads to a pro-fibrotic environment.

The distinct association of alveolar bronchiolization with fibrotic regions of the lung led us to initially hypothesize that ectopic KRT5+ basal cells would directly drive a fibrotic fibroblast transition. Instead, we observed induction of the inflammatory state upon CM treatment. The *in* vivo association with dysplastic epithelium and fibrotic remodeling therefore cannot be explained by direct fibrotic fibroblast conversion and instead implicates a multi-step process by which the accumulation of inflammatory fibroblasts indirectly promotes tissue remodeling due to persistent recruitment of inflammatory infiltrate. Indeed, chronic immune activation is well established to contribute to tissue damage and fibrosis, highlighted by research into the COVID-19 induced cytokine storm that received significant attention in recent years (*59–64*). Further supporting this idea, ectopic KRT5+ basal cells have recently been directly implicated in the recruitment of CD4+ effector and CD8+ T Cells to sites of alveolar bronchiolization (*19*). This inhibition of euplastic repair by T-cell derived inflammatory cytokines likely creates further opportunity for fibrotic remodeling (*65*).

IL-1α and IL-1β are both critical pro-inflammatory cytokines that signal through IL1R1, but vary greatly in their regulation and expression by cell type. IL-1β is produced primarily by immune cells and released in response to external stimuli, whereas many non-immune cell types, especially basal keratinocytes, are known to constitutively express IL-1α (*25*, *42*, *43*, *66*, *67*). Both are translated as precursor proteins, where inflammasome / caspase-dependent processing is required to produce the mature, bioactive form of IL-1β and enhance the activity of the already bioactive pro-IL-1α. IL-1α is thought to play a more immediate role in local inflammation, whereas IL-1β is thought to possess more systemic inflammatory functions (*41*). IL-1α released from epithelial cells undergoing necrotic cell death is a well-appreciated early warning signal during infection or injury (*42*, *68*). While the release of IL-1α from dying cells is undoubtedly one source of the cytokine during lung injury, regions of alveolar bronchiolization are chronic pathologic features that persist long after injury resolution. Ectopic KRT5+ cells thus represent active, long-lived sources of the inflammatory cytokine IL-1α in regions of alveolar bronchiolization, capable of driving nearby fibroblasts into a pro-inflammatory phenotype and propagating cellular pathology across multiple lineages.

The inflammatory fibroblast phenotype has been identified in multiple studies and their gene expression profile suggests their likely involvement in immune cell recruitment. Indeed, after bleomycin treatment, mice with alveolar fibroblasts lacking TGFβR2 experienced significantly greater monocyte/myeloid recruitment to the lungs, coincident with a significant enrichment in the inflammatory fibroblast population and increased mortality during the inflammatory phase of the injury (*31*). Inflammatory fibroblasts have additionally also been proposed to act as an intermediate state that alveolar fibroblasts pass through prior to the assumption of a fibrotic phenotype (*31*, *69*). However, our TGF-β titration studies indicate that IL-1α-driven inflammatory fibroblasts do not appear primed to assume the fibrotic phenotype (Supplemental Figure 6), at least as judged by sensitivity to TGF-β. While the inflammatory fibroblast phenotype may still represent an intermediate state, this does not apparently entail increased responsiveness to TGF-β. Interestingly, we found that culture on a stiffer substrate alone appeared to promote similar immune-related processes as CM treatment, suggesting that fibroblasts in fibrotic / mechanically stiffer regions of the lung may be prone to assuming the inflammatory phenotype (Supplemental Figure 7).

Many cytokines and chemokines were significantly upregulated in the inflammatory fibroblast population, but CCL2 was amongst the most highly upregulated. In fact, a number of other chemokines important for monocyte recruitment were also highly upregulated, including *Ccl3, Ccl7*, and *Ccl20*. This is particularly notable given that CCL2 and CCR2+ cells are recognized to contribute to fibrotic disease across multiple organs by modulating the recruitment and activation of immune cells (*70–72*). CCR2+ cells have been found to promote a sustained inflammatory environment through the persistent recruitment of neutrophils, and promote fibrosis through aberrant TGF-β signaling (*73*). The strong trend toward a reduction in CCR2+ cells upon IL-1α neutralization alongside the significant reduction in inflammatory fibroblasts implicate a mechanism by which ectopic KRT5+ basal cells indirectly drive PF through the induction of persistent inflammatory fibroblasts and subsequent sustained recruitment of CCR2+ immune cells to sites of alveolar bronchiolization.

Our studies indicate that rather than promoting the conversion of nearby fibroblasts to a fibrotic phenotype, dysplastic basal cells reprogram fibroblasts into an inflammatory state, driving aberrant immune cell recruitment and chronic inflammation. As such, dysplastic basal cells may “set the stage” for fibrotic remodeling without directly producing the pro-fibrotic signals ultimately associated with pathologic extracellular matrix deposition. Indeed, an indirect role for dysplastic basal cells in promoting PF could explain their consistent association with fibrotic regions without necessitating direct induction of a fibrotic fibroblast phenotype. Taken together, the long-term alveolar persistence of these dysplastic cells, their known role in T cell recruitment, and their indirect role in monocyte recruitment all represent major contributors to their capacity to promote chronic pathology.

## Methods

### Study Design

In this study, we investigated signaling between ectopic KRT5⁺ basal cells and surrounding fibroblasts after influenza-induced lung injury. To this end, we employed spatial transcriptomics, bulk RNA-sequencing, lineage tracing, microscopy, controlled cell culture perturbation experiments, and delivered neutralizing antibodies *in vivo*. Experiments were performed using both influenza-infected mice and primary cell cultures derived from mouse lungs.

Sample sizes varied by experimental approach and are explicitly described in figure legends. For *in vivo* experiments, we aimed to include ≥4 mice per group, but group sizes were sometimes limited by mortality, insufficient disease severity, or genotype availability. No statistical methods were used to predetermine sample sizes, but sample sizes were consistent with those reported in prior studies. Rules for stopping data collection were defined in advance: mice experiencing > 30% weight loss were euthanized, and final endpoints were defined as specific days post–influenza infection, depending on the experiment. Inclusion criteria for influenza experiments required >10% weight loss and mice with <10% weight loss were excluded. Experiments were independently repeated three times, with one to four technical replicates per experiment. Animals were randomly assigned to experimental groups at the start of each experiment, with stratification to balance sex across groups. All mice were obtained from the same source population and age range. Investigators were not blinded to group allocation. Specific sample sizes, numbers of replicates, and statistical tests are provided in the figure legends.

### Animals and Treatment

In this study, unless otherwise specified, adult 8 to 16-week-old mice were used for all experiments. Males and females were assigned in approximately equal proportions and experimenters were not blinded to mouse age or sex. All animal experiments were reviewed and approved by the University of Pennsylvania’s Institutional Animal Care and Use Committee (IACUC) and followed all National Institutes of Health (NIH) Office of Laboratory Animal Welfare regulations, protocol 806262. The mice used in the experiments were either wild-type (C57BL/6J) or crosses between the following transgenic mice: Ai14tdTomato (Gt(ROSA)26Sortm14(CAG-tdTomato)Hze), Krt5CreERT2, PDGFRα-H2B:EGFP, and Scube2CreERT2 (*31*, *36*, *37*, *74*). Genotyping primers for all transgenic mice used are as follows: Ai14tdTomato WT-Forward: AAGGGAGCTGCAGTGGAGTA, Ai14tdTomato WT-Reverse: CCGAAAATCTGTGGGAAGTC, Ai14tdTomato MUT-Forward: CTGTTCCTGTACGGCATGG, Ai14tdTomato MUT-Reverse: GGCATTAAAGCAGCGTATCC, Krt5CreERT2 Common-Forward: GCAAGACCCTGGTCCTCAC, Krt5CreERT2 WT-Reverse: GGAGGAAGTCAGAACCAGGAC, Krt5CreERT2 MUT-Reverse: ACCGGCCTTATTCCAAGC, PDGFRα-H2B:EGFP WT-Forward: CCCTTGTGGTCATGCCAAAC, PDGFRα-H2B:EGFP WT-Reverse: GCTTTTGCCTCCATTACACTGG, PDGFRα-H2B:EGFP Mut-Reverse: ACGAAGTTATTAGGTCCCTCGAC, Scube2CreERT2 Common-Forward: GAGTCCCGAGAGATGTTTCC, Scube2CreERT2 WT-Reverse: ACAGTGCTACTCAAAGCGT, and Scube2CreERT2 MUT-Reverse: GTCCATCAGGTTCTTGCGA.

### Influenza-induced Lung Injury

Mice were housed in Animal Biosafety Level 2 (ABSL-2) facilities for all influenza experiments. For infection, mice were first anesthetized using an anesthesia vaporizer system set at 3.5% isoflurane (Penn Veterinary Supply, Inc.; VED1350) in 100% O2 and agonal breathing was visually confirmed. Afterwards, influenza virus A/H1N1/PR/8 was delivered intranasally with respect to mouse weight (0.7 * bodyweight in grams) TCID50 PR8 (ex: A 20g mouse received (0.7 * 20g) = 14U PR8) dissolved in 30uL of phosphate-buffered saline (PBS; Corning MT21-031-CM) as previously described. Control mice were administered the same volume of PBS. Infected mice that lost ≥10% of their starting body weight by day 9 post flu were considered adequately infected and used for all experiments involving influenza infection. Mice were regularly weighed throughout infection and euthanized at the endpoints specified for harvest.

### Tamoxifen Administration and Neutralizing Antibody Delivery

For intraperitoneal (IP) tamoxifen experiments, tamoxifen (Toronto Research Chemicals, T00600) was dissolved in 50uL of corn oil and administered through injection (250 mg kg−1 body weight) into the intraperitoneal cavity. For oral gavage tamoxifen experiments (Scube2-CreERT2; PDGFRα-H2B:EGFP; Ai14tdTomato mice), we administered Tamoxifen (200mg/kg) via oral gavage on 3 consecutive days 2 weeks prior to infection. Mice homozygous for Scube2-CreERT2 were used. For IP antibody experiments, both InVivoMAb anti-mouse IL-1α (Bioxcell, BE0243; Clone ALF-161) and InVivoMAb polyclonal Armenian hamster IgG (Bioxcell, BE0091; Polyclonal) were resuspended in PBS at 2μg/μL and 100μL was delivered through IP injection. Control mice for all injections received the corresponding volume of vehicle control unless otherwise stated.

### Lung Tissue Preparation and OCT Embedding

Following Euthanasia, the heart was thoroughly perfused with cold PBS via the left atrium, and the lungs were subsequently washed repeatedly with PBS. The lungs were inflated with 1 mL 3.2% paraformaldehyde (PFA), tied off, and fixed at room temperature (RT) for 1 hour. Tissue was then washed multiple times in PBS at RT over 1 hour, followed by an overnight incubation in 30% sucrose (Millipore Sigma, S8501) in PBS with 0.02% sodium azide (Millipore Sigma, S2002) at 4 °C while rocking. Lungs were then incubated in 15% sucrose in PBS with 0.02% sodium azide and 50% optimal cutting temperature (OCT) compound (Fisher Scientific, 23-730-571) with gentle agitation at RT for 2 hours. Lung lobes were randomly oriented in molds and embedded in OCT by flash freezing in a dry ice-ethanol bath.

### Sectioning and Immunostaining

7-μm-thick sections were cut with the Leica CM3050 S Research Cryostat (Leica Biosystems) and stored at −20°C. Sections were taken from multiple discrete (>100μm) distant) levels of each block to survey a significant portion of the lung. Tissue sections were brought to RT, post-fixed for 5 minutes in 3.2% PFA, rinsed with PBS, and blocked at RT for 1 hour in blocking buffer (2% BSA (GoldBio, A-420-10), 5% donkey serum (Millipore Sigma, S30-100mL) and 0.1% Triton X-100 (FisherScientific, BP151-100) in PBS). Slides were then incubated in primary antibody (detailed below) diluted in blocking buffer at 4°C overnight. The next day, slides were rinsed with PBS + 0.1% Tween 20 (PBST; Sigma-Aldritch, 9005-64-5) and incubated with fluorophore-conjugated secondary antibodies (detailed below) diluted 1:1000 in blocking buffer at RT for 1 hour. Slides were washed once more with PBST, incubated in 1μM DAPI (Fisher Scientific, D1306) for 5 minutes, rinsed with PBS, and mounted with either Fluoroshield (Millipore Sigma, F6182) or Prolong Gold (ThermoFisher, P36930) overnight. Slides stained for CCR2 were additionally treated with TrueBlack® Plus Lipofuscin Autofluorescence Quencher (Biotium, 23014) to reduce background fluorescence. Primary antibodies: Chicken anti-KRT5 (1:1000, BioLegend, clone poly 9059, 905901), Rabbit anti-KRT5 (1:1000, BioLegend, polyclonal, 905504), Rabbit anti-COL1A1 (1:500, Monoclonal E8F4L, Cell Signalling, 72026), SAA3 Rat anti-Mouse Alexa Fluor™ 647 (1:200, BD Pharmingen™, Clone: JOR110A, 566652), and PE anti-mouse CCR2 (1:300, Clone: SA203G1, BioLegend, 150609). Secondary antibodies: Donkey anti-Rat IgG (H+L), Alexa Fluor™ 568 (Invitrogen, A78946), Donkey anti-Rat IgG (H+L), Alexa Fluor™ 647 (Invitrogen, A78947), Donkey anti-Chicken IgY (H+L), Alexa Fluor™ 488 (Invitrogen, A78948), Donkey anti-Rabbit IgG (H+L), Alexa Fluor™ 647 (Invitrogen, A31573), Donkey anti-Rabbit IgG (H+L), Alexa Fluor™ 488 (Invitrogen, A21206), and Donkey anti-Rabbit IgG (H+L), Alexa Fluor™ 568 (Invitrogen, A10042).

For alveolar fibroblast lineage-trace experiments, following fixation, lungs were washed three times in ice-cold PBS, sectioned into three equal transverse pieces, and embedded in 4% low-melting-point agarose (IB70050, IBI Scientific). Thick-cut lung slices (TCLS) were generated at 150 µm thickness using a Leica VT1000S vibratome. Sections were permeabilized in PBS containing 1% Triton X-100 for 1 hour at room temperature and incubated with primary antibodies diluted in blocking buffer (PBS containing 0.3% Triton X-100 and 1% bovine serum albumin; B2518, Sigma-Aldrich) for 48 hours at 4°C with gentle rocking. Sections were washed three times in permeabilization buffer (30 min per wash at room temperature) and incubated with secondary antibodies diluted in blocking buffer for 48 hours at 4 °C with gentle rocking. Following immunostaining, sections were cleared with Scale A2 for one week before imaging by confocal microscopy as described previously (*75*). For confocal imaging, primary antibodies used were: chicken anti-GFP (1:500; GFP-1020, Aves Labs), guinea pig anti-RFP (1:500; 390004, Synaptic Systems), rabbit anti–keratin 5 (1:200; 905501, BioLegend), and rat anti-SAA3 (1:300; ab231680, Abcam). Secondary antibodies were used at 1:500 dilution and included: donkey anti-rabbit IgG Alexa Fluor 405 (A48258, Invitrogen), donkey anti-rat IgG Alexa Fluor 647 (A48272TR, Invitrogen), Cy3-conjugated donkey anti–guinea pig IgG (H+L) (706-165-148, Jackson ImmunoResearch), and donkey anti-chicken IgG Alexa Fluor 488 (703-545-155, Jackson ImmunoResearch).

### Microscopy

For 7-µm sections, images were taken on a Leica DMi8 microscope with a Leica DFC9000 sCMOS camera using Leica Application Suite X (LAS X) software. Immunofluorescent and phase contrast images were taken at 20x or 63x using z-stacks to perform the extended depth-of-field function. One to five 20x images were taken of KRT5 regions found on each stained slide.

For confocal imaging, images were acquired using a Leica SP8 confocal microscope and processed using ImageJ (FIJI). TCLS images were acquired as z-stacks spanning 100–150 µm with optical sections collected at 2 µm intervals using a 40x objective. Final images consisted of merged tile scans generated from five z-stack projections.

### Quantification of Immunofluorescence Data

Images taken during the same microscope session were subjected to identical adjustments for uniform comparison and analysis using ImageJ. Thresholding was performed at the same value to quantify the area of both SAA3 and KRT5 signal, while CCR2+ cells were counted manually.

For TCLS, three-dimensional image analysis was performed using Imaris software (Oxford Instruments). The Surfaces function was applied to the keratin 5 channel to generate a 3D keratin 5 pod surface. Next, the Colocalization analysis tool was used to generate a new channel for PDGFRα-GFP+/Scube2-lin+ fibroblasts. Similarly, the tool was used to find and build channels for SAA3+/PDGFRα-GFP+ and SAA3+/PDGFRα-GFP+/SCUBE2-lin+ fibroblasts. Subsequently, all channels were masked to the 3D keratin 5 pod surface using the Surface Masking tool. Quantification of PDGFRα-GFP⁺, PDGFRα-GFP⁺/SCUBE2-lin⁺, SAA3⁺/PDGFRα-GFP⁺, and SAA3⁺/PDGFRα-GFP⁺/SCUBE2-lin⁺ fibroblasts was performed using the Spot Detection tool (spot diameter = 7 µm; quality threshold > 4.25).

### Lung Tissue Preparation for Primary Cell Culture

Following euthanasia the heart was perfused and the lungs were washed with PBS as described above for OCT embedding. Lung cells were isolated by inflating the lungs with digestion buffer consisting of 15 U/mL dispase II (Gibco, 17105041) in Hank’s Balanced Salt Solution (HBSS; Corning, MT21-023-CV) and 1:1000 deoxyribonuclease I (DNASE I; Millipore Sigma, D4527), tying them off, and incubating for two minutes before separating the lobes from the mainstem bronchi. The lobes were then minced and incubated in digestion buffer for 45 minutes while shaking at RT. The digested lung was then further mechanically dissociated by pipetting in DMEM (Gibco, 11965092) + 10% cosmic calf (CC) serum +2% penicillin-streptomycin (PS) and passed through a 70 μM filter before being pelleted and resuspended in ACK Lysis buffer for 5 minutes to digest red blood cells. The cells were then spun down, resuspended in DMEM + 2% CC +2% PS, and incubated at 37°C for 30 minutes to recover. Finally, the cells were pelleted, washed with PBS, and resuspended as a single cell suspension in media corresponding to the cell type to be cultured.

### Ectopic Basal Cell Culture and Passaging

A single cell suspension of 21-day post-infection lungs was prepared as described above, resuspended in PneumaCult™-Ex Plus Medium (Stemcell Technologies, 05040) + 1x PneumaCult™-Ex Plus Supplement (Stemcell Technologies, 05040) + 1:1000 hydrocortisone stock solution (Stemcell Technologies, 07925) + 1μM A8301 (TGF-β inhibitor, Millipore Sigma, SML0788) + 10μM Y-27632 (ROCK inhibitor, Cayman Chemical, 10005583) + 1:5000 Primocin® (Invivogen, ant-pm-05), and plated across two 10cm dishes. The cells were allowed to grow until 80% confluent with media changes every 48 hours. Basal cells expanded in these 10cm dishes were frozen down and new experiments began with a thawed vial of cells below passage 6. To passage, cells were treated with 0.05% trypsin-EDTA (ThermoFisher, 25200056) in PBS for 10 min at 37°C, quenched with DMEM + 10% CC + 1% PS, and pelleted at 550 × g for 5 minutes at 4°C.

### Pulmonary Fibroblast Culture and Passaging

A single cell suspension of uninjured lungs was prepared as described above, resuspended in DMEM + 10% CC + 1% PS, and plated on a single 10cm dish. The cells were allowed to reach 90% confluency with media changes every 48 hours and a reduction to 2% CC on day 4 of expansion. The fibroblasts were then frozen down, and new experiments began with freshly thawed vials of passage 1 cells in DMEM + 2% CC + 1% PS. Fibroblasts were cultured on plastic and either 0.5kPa or 16kPa Cytosoft™ (Advanced Biomatrix, 5140(0.5kPa), 5143 (16 kPa)) plates. Cytosoft™ 6-well plates were pre-incubated with 1:30 PureCol1 (Advanced Biomatrix, 5005) in PBS for 1 hour at RT, washed with 5mL PBS, then 5mL culture media prior to plating cells.

### Conditioned Media Generation

To generate conditioned media for cell culture experiments, basal cells were expanded on 10cm dishes until reaching 90% confluency. The media was then aspirated, the dishes were washed twice with 10mL PBS, and 6mL of DMEM + 2% CC + 1% PS was added to each 10cm dish. The cells were allowed to culture for 24 hours, conditioning the media, after which the supernatant was transferred to a conical and cells were pelleted at 1000 x g for 10 minutes at 4°C. The media was carefully decanted without disturbing the cell pellet. For larger batches of conditioned media, multiple separate dishes were combined, re-aliquoted, then stored at −80°C until being thawed immediately prior to use. Conditioned media for ELISA purposes was prepared as described above but was not put through a freeze-thaw cycle. Fibroblast conditioned media was generated from passage 1 naïve unstimulated, or Conditioned Media stimulated, fibroblasts expanded in a 10cm dish until reaching confluency.

### Media Supplementation and Depletion Experiments

Recombinant proteins, inhibitors, antagonists, and neutralizing antibodies used in these experiments were added directly to the respective culture media as noted in figure legends, while control media received only vehicle. The working concentration for the following reagents can be found in the text and figure legends: Human IL-1RA Recombinant Protein (IL1Ra) (PeproTech®, 200-01RA), Mouse IL-1 alpha Recombinant Protein, (PeproTech®, 211-11A), Mouse IL-1α Neutralizing Antibody (Invivogen, Anti-mIL-1α-mIgG1 clone 6H7, mIL-1α-mab9-02), Recombinant Mouse IgG1 Isotype Control Antibody (Invivogen, Anti-β-Gal-mIgG1 clone T9C6, bgal-mab9-02), Human TGF-β 1 Recombinant Protein (PeproTech®, 100-21), Human GDF15 Recombinant Protein (PeproTech®, 120-28C), Human VEGF Recombinant Protein (PeproTech®, 450-32), Human OPN Recombinant Protein (PeproTech®, 120-35), Murine CXCL1 Recombinant Protein (PeproTech®, 250-11), Human IGFBP3 Recombinant Protein (PeproTech®, 100-08), Human TNFRSF11B Recombinant Protein (PeproTech®, 450-14), Murine CXCL5 Recombinant Protein (PeproTech®, 250-17), and Muring CCL20 Recombinant Protein (PeproTech®, 250-27).

For IL-1α antibody depletion experiments, 100ng/mL of Mouse IL-1α Neutralizing Antibody or IgG1 Isotype Control Antibody was added to freshly generated conditioned media and incubated at 4°C while rocking for 30 minutes, followed by the addition of excess (1:200) TrueBlot® Anti-Mouse IgG Magnetic Beads (Rockland, Goat Polyclonal, 00-1811-20) and subsequent incubation at room temperature while rocking for 30 minutes. The magnetic beads, antibody, and bound protein was then depleted by passing the media through LS Columns (Miltenyi Biotec, 130-042-401) on a QuadroMACS™ Separator (Miltenyi Biotec, 130-090-976). Media was then stored at −80°C and thawed immediately prior to use.

### IL-1α ELISA

Freshly generated conditioned media from naïve unstimulated fibroblasts and from basal cells was used to quantify secreted IL-1α with the ELISA MAX™ Deluxe Set Mouse IL-1α (BioLegend; 433404) according to their protocol.

### Bulk RNA-seq Experiments

For fibroblast bulk RNA-sequencing culture experiments, four technical replicates per group were plated on either plastic or 0.5 kPa cytosoft plates and cultured in either control or conditioned media. Cells were initially plated in DMEM + 2% CC + 1% PS and allowed to grow overnight. After 24 hours, media was replaced with control or conditioned media and cells were treated for a total of 48 hours, with a media refresh at 24 hours. RNA was extracted from all samples as described in “RNA Isolation and RT-qPCR” and shipped on dry ice to Genewiz (Azenta Life Sciences) for library preparation and sequencing. PolyA-selected RNA was used to generate libraries using the NEBNext Ultra II RNA Library Prep Kit for Illumina (NEB) according to the manufacturer’s instructions. Sequencing was performed using a 2 × 150bp paired-end configuration on an Illumnia HiSeq at a read depth of ∼45 million reads per sample on average. Sequencing reads were aligned to the GRCm39 mouse reference genome using Kallisto and imported into R (RStudio) using the tximport package. Transcripts were excluded from downstream analysis if they had zero counts across any sample or failed to reach a minimum expression threshold of one count per million in at least four samples. Data was then normalized using the trimmed mean of M values method using the edgeR package. Voom was then used to model the mean-variance relationship, followed by linear modeling and empirical Bayes moderation using the ImFit and eBayes functions from the Limma package. This yielded differentially expressed genes between the four groups: Plastic control, Plastic CM, 0.5 kPa control, and 0.5 kPa CM.

### RNA Isolation and RT-qPCR

RNA was isolated using the either the ReliaPrep™ RNA Cell Miniprep kit (Promega, Z6011) or Direct-zol RNA Miniprep Plus (ZymoResearch; R2072) and concentration/quality was assessed via nanodrop. RNA >100ng was amplified to generate cDNA libraries with iScript™ Reverse Transcription Supermix (BioRad, 1708841) and a dilution of amplified cDNA in DI water was used as input in each well for RT-qPCR. RT-qPCR was performed on the Applied Biosystems QuantStudio 6 Real-Time PCR System (ThermoFisher) with PowerUp SYBR Green Master Mix (Applied Biosystems, A25742). Gene expression was calculated (ΔΔCt method) relative to housekeeping gene Rpl19 within each sample and plotted as fold change over average control expression for each gene. The following primers were used for RT-qPCR: Rpl19-Forward: ATGAGTATGCTCAGGCTACAGA, Rpl19-Reverse: GCATTGGCGATTTCATTGGTC, Ccl2-Forward: TTAAAAACCTGGATCGGAACCAA, Ccl2-Reverse: GCATTAGCTTCAGATTTACGGGT, Saa3-Forward: TGCCATCATTCTTTGCATCTTGA, Saa3-Reverse: CCGTGAACTTCTGAACAGCCT.

### Spatial Transcriptomics

Krt5-CreERT2/ Ai14tdTomato mice were infected with the flu, injected with IP tamoxifen at day 11, and lungs were harvested at day 21, embedded in OCT, and sectioned onto poly-L-lysine coated glass slides (Sigma-Aldrich, P0425). Sample RNA quality was assessed by harvesting RNA from sections using RNeasy FFPE Kit (Qiagen, 73504) and the lungs with the highest quality RNA were further sectioned and a lineage-positive lobe was selected to perform VISIUM Spatial Transcriptomics alongside a lobe from a healthy mock infected mouse. The selected slides were delivered to the Single Cell Technology Core at The Children’s Hospital of Philadelphia for H&E staining / imaging and further processing. Immediately after imaging, the coverslips were removed and the tissues were processed for destaining and decrosslinking, followed by probe hybridization, ligation, release, extension, pre-amplification, and probe-based library construction, using the Visium CytAssist Spatial Gene Expression for FFPE, Mouse Transcriptome, 6.5 mm kit (PN-1000521), along with the Visium Mouse Transcriptome Probe Set v1.0 (PN-1000365) and the Dual Index Plate TS Set A (PN-1000251), following the protocols outlined in CG000662 and CG000495. Library pooling and sequencing were handled by the CHOP HTS Core. For details about the sequencer, flow cell used, and actual sequencing depth, please contact Teodora. Recommended sequencing parameters can be found in the Visium CytAssist Spatial Gene Expression Reagent Kits User Guide, pages 103-104. Raw 10x Genomics Visium data was run through the Space Ranger pipeline without further modification and aligned to the mm10 reference genome. Images and figures were generated using Loupe Browser (10x Genomics).

### Cytokine Profiling

Two replicates of freshly generated conditioned media and control media were used to probe for the presence and relative quantity of 111 selected cytokines and chemokines using the Proteome Profiler Mouse XL Cytokine Array (R&D Systems, ARY028). The kit was used according to the manufacturers protocol, analyzed using a ChemiDoc MP Imaging System (Bio-Rad) and Image Lab software (Bio-Rad); Images were equally adjusted to ensure direct qualitative comparison.

## Statistical Analysis

Graphs were made and statistical analysis was performed with either GraphPad Prism 10 or on RStudio. Data was pooled from independent experiments for analysis and the statistical tests used for each experiment can be found in their respective figure legends. A *P*-value less than 0.05 was considered significant in this study (\**P* < 0.05, \*\**P* < 0.01, \*\*\**P* < 0.001, \*\*\*\**P* < 0.0001, ns: not significant).

## Acknowledgements

We thank all members of the Vaughan and Zepp labs for discussions; the Single Cell Technology Core at The Children’s Hospital of Philadelphia for their assistance with the VISIUM spatial gene expression experiments; Dr. Dean Sheppard for providing us with the Scube2-CreERT2 mice. We are grateful for the generous support provided by the Ayla Gunner Prushansky Research Fund.

## Author Contributions

N.P.H. designed experiments, performed the majority of experiments, acquired and analyzed data, prepared figures, and wrote the manuscript. R.Z.S. performed experiments, acquired and analyzed data, prepared figures, and contributed to manuscript writing. A.K., J.W., S.K., M.Me., M.M.M., E.A.M., D.M.A. performed experiments, contributed to scientific discussion and interpretation of results. M.S. and H.K. performed genotyping and contributed to scientific discussion. J.A.Z. and A.E.V. conceived and oversaw the study, supervised experimental design, and oversaw manuscript preparation. All authors reviewed and approved the final manuscript.

## Data & Material Availability

All data required to evaluate the conclusions made in this manuscript are present in the paper or the supplementary materials. The murine bulk RNA-seq raw and processed data files in this study has been deposited in the Gene Expression Omnibus (GEO) under accession number GSE326787. The murine_VISIUM raw and processed data files have been deposited in the GEO under accession number GSE326788.

---

**Supplemental Figure 1:**
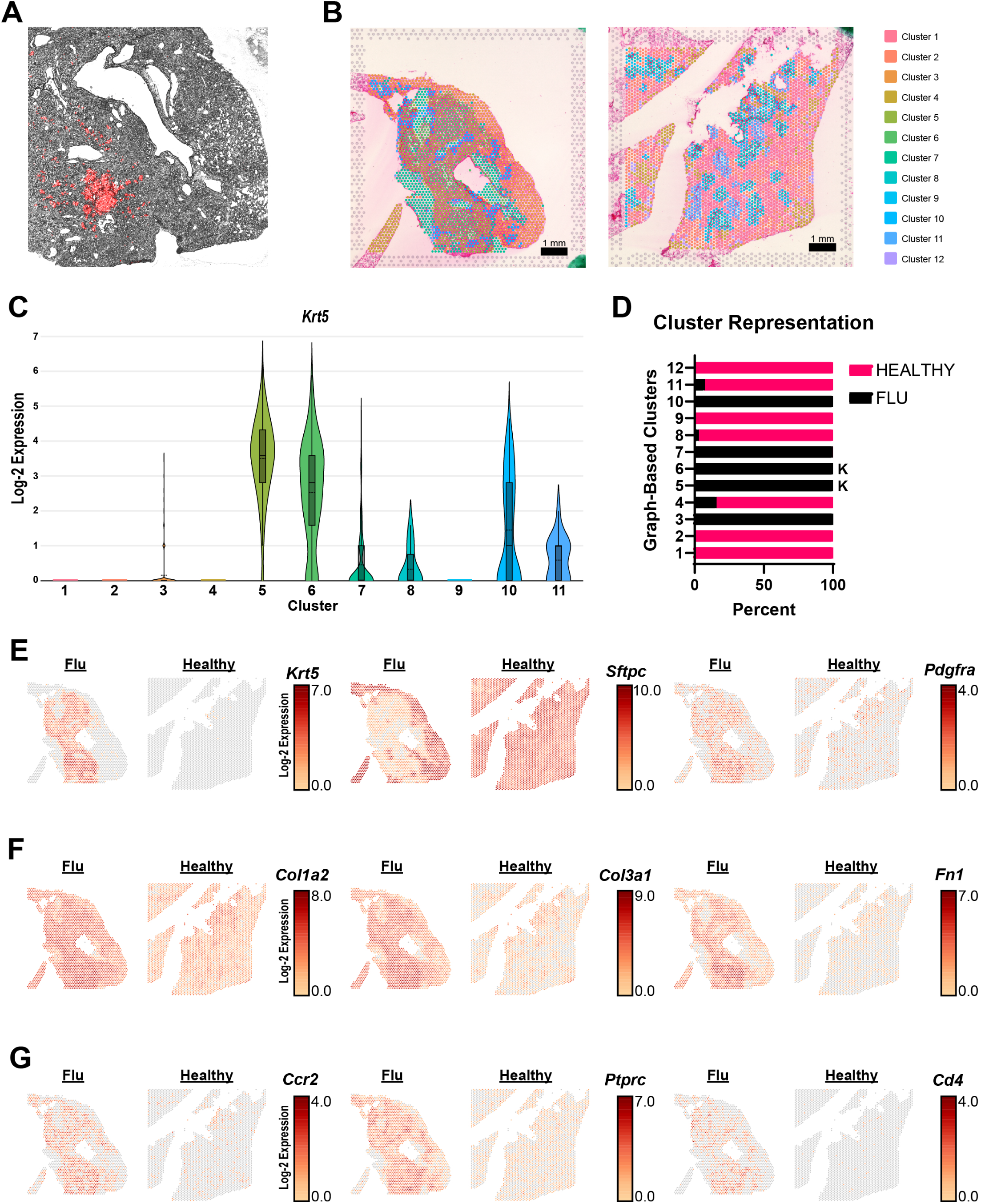
Inflammatory and Fibrotic Transcriptional Programs are Enriched within Regions of Alveolar Bronchiolization. **(A)** Phase contrast image overlaid with KRT5-CreERT2 lineage-trace (RED) of the post-flu lung selected for spatial transcriptomic analysis. **(B)** Graph-based clustering of both the post-flu (left) and uninjured, healthy (right) lung superimposed over the respective H&E images. **(C)** Log-2 *Krt5* expression plotted as a violin plot for the graph-based clusters. **(D)** The percentage of graph-based clusters that belong to either the healthy or post-flu lung. The “*K*” indicates the two clusters containing the highest *Krt5* expression. **(E)** Spatial sequencing plots of the post-flu lung (left) and healthy lung (right) showing the log-2 expression for *Krt5*, *Sftpc*, and *Pdgfra*. **(F)** Spatial sequencing plots of the post-flu lung (left) and healthy lung (right) showing the log-2 expression of fibrotic marker genes *Col1a2*, *Col3a1*, and *Fn1*. **(G)** Spatial sequencing plots of the post-flu lung (left) and healthy lung (right) showing the log-2 expression of inflammatory marker genes *Ccr2*, *Ptprc* (*Cd45*), and *Cd4*.

**Supplemental Figure 2:**
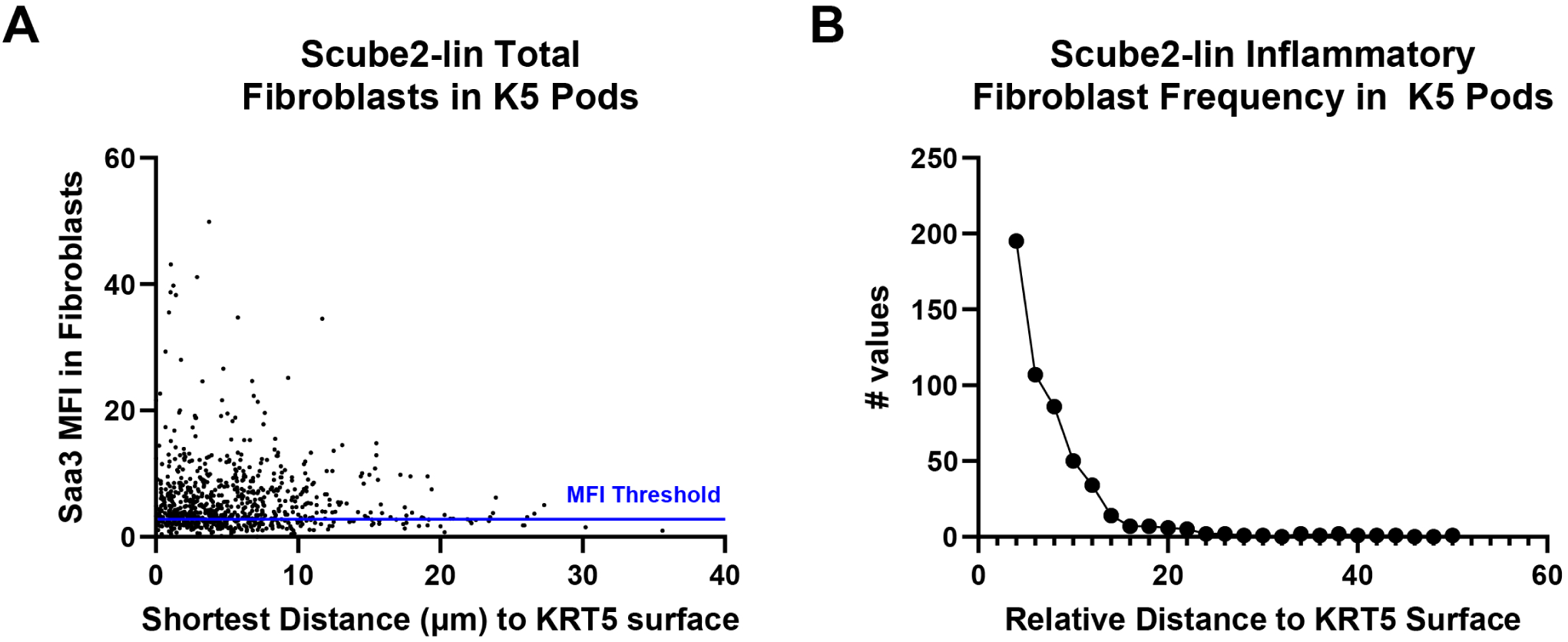
Dysplastic KRT5+ Regions Exhibit a Proximity-Dependent Enrichment of the Inflammatory Fibroblast Population. **(A)** Scatterplot summarizing the total number of Scube2-lineage-positive fibroblasts and their corresponding mean fluorescent intensity (MFI) of SAA3 as a function of distance to KRT5+ regions *in vivo*. Blue line at 3.0 MFI indicates the threshold at which a cell is determined to SAA3 positive. **(B)** Frequency distribution of inflammatory fibroblasts (SAA3 MFI ≥ 3) in (A).

**Supplemental Figure 3:**
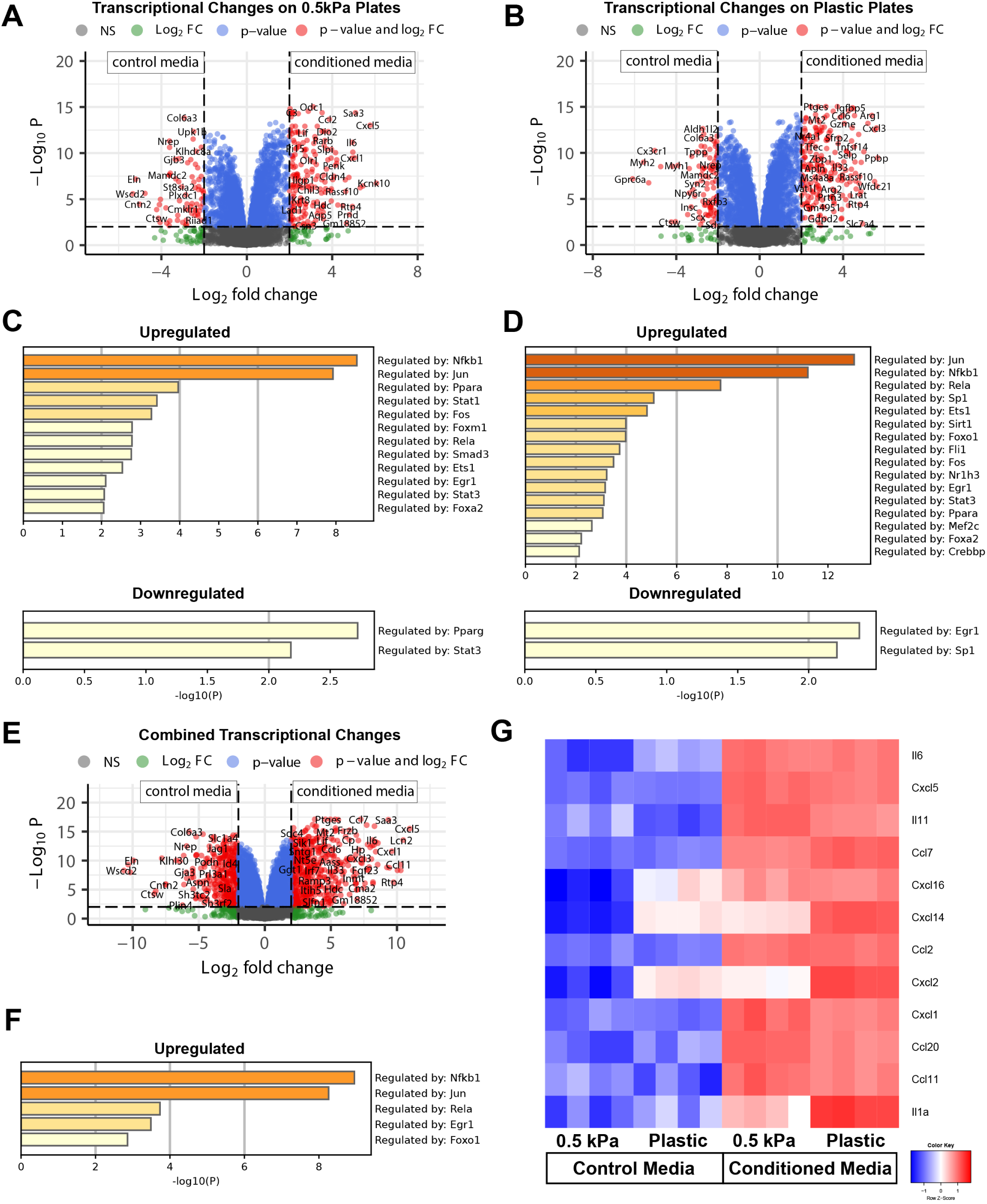
Fibroblasts Exhibit Robust Transcriptomic Changes Upon Exposure to Dysplastic Basal Cell-Derived Factors. **(A)** Volcano plot illustrating significantly differentially expressed genes between control and conditioned media treated fibroblasts cultured on 0.5kPa plates with genes labeled. *P* value < 0.01, Fold Change >4,184 upregulated 68 downregulated. **(B)** Volcano plot illustrating significantly differentially expressed genes between control and conditioned media treated fibroblasts cultured on plastic plates with genes labeled. *P* value < 0.01, Fold Change >4, 203 upregulated 94 downregulated. **(C)** Enrichment analysis of transcription factors (TRRUST database via Metascape) using the significantly up and down-regulated genes in (A). **(D)** Enrichment analysis of transcription factors (TRRUST database via Metascape) using the significantly up and down-regulated genes in (B). **(E)** Volcano plot illustrating significantly differentially expressed genes between control and conditioned media treated fibroblasts cultured on both 0.5kPa and plastic plates with genes labeled. *P* value < 0.01, Fold Change >4, 538 upregulated 382 downregulated. **(F)** Enrichment analysis of transcription factors (TRRUST database via Metascape) using the top 92 significantly up-regulated genes (*P* value < 0.01, Fold Change >4) shared between 0.5kPa and plastic conditioned media treated fibroblasts. **(G)** Heatmap depicting the relative expression of a variety of inflammatory marker genes upregulated in conditioned media treated fibroblasts.

**Supplemental Figure 4:**
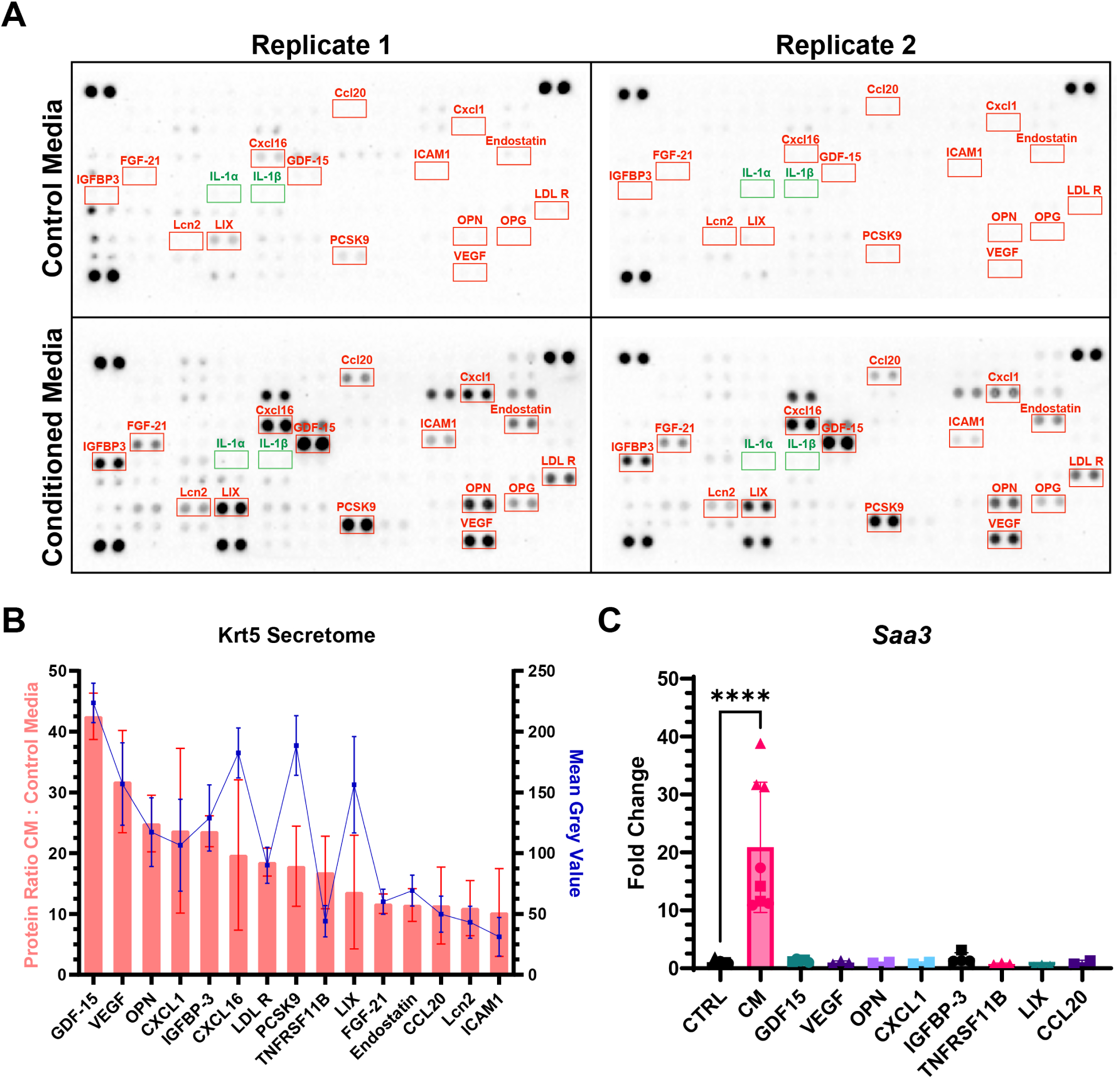
Proteome Profiler Mouse Cytokine Array Highlights Relative Abundance of Secreted Proteins in Conditioned Media. **(A)** Scan of the two proteome profiler blots with the top 15 proteins enriched in the conditioned media relative to control media highlighted in red. Note that both IL-1α and IL-1β (highlighted in green) were undetectable. **(B)** Plot showing the relative protein ratio and mean grey value of the enriched proteins in (A). **(C)** Recombinant proteins supplemented in control media failed to induce a significant upregulation of *Saa3* in fibroblasts cultured on plastic. Recombinant proteins were supplemented at the following concentrations: GDF15 (200ng/mL), VEGF (10ng/mL), OPN (200ng/mL), CXCL1 (100ng/mL), IGFBP3 (200ng/mL), TNFRSF11B (10ng/mL), CXCL5 (100ng/mL), and CCL20 (10ng/mL). (C) *P* values were calculated using a one-way ANOVA with post-hoc Turkey multiple comparison test: \*\*\*\**P* < 0.0001.

**Supplemental Figure 5:**
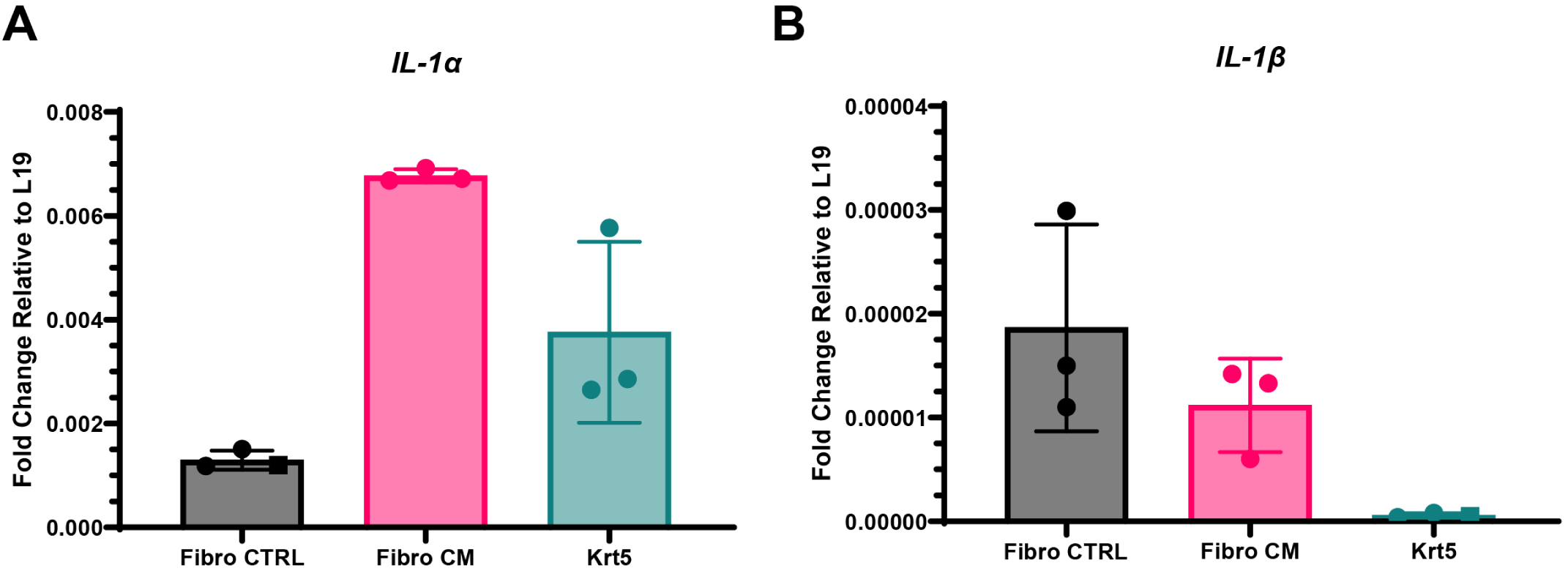
Relative IL1 Transcripts in Treated and Untreated Fibroblasts and Dysplastic Basal Cells. **(A)** *IL1a* **(B)** *IL1b* gene expression by RT-qPCR for fibroblasts cultured on plastic before (Fibro CTRL) and after (Fibro CM) conditioned media treatment as well as the KRT5+ Cells used to generate the same conditioned media. Fold change is plotted relative to housekeeping gene Rpl19.

**Supplemental Figure 6:**
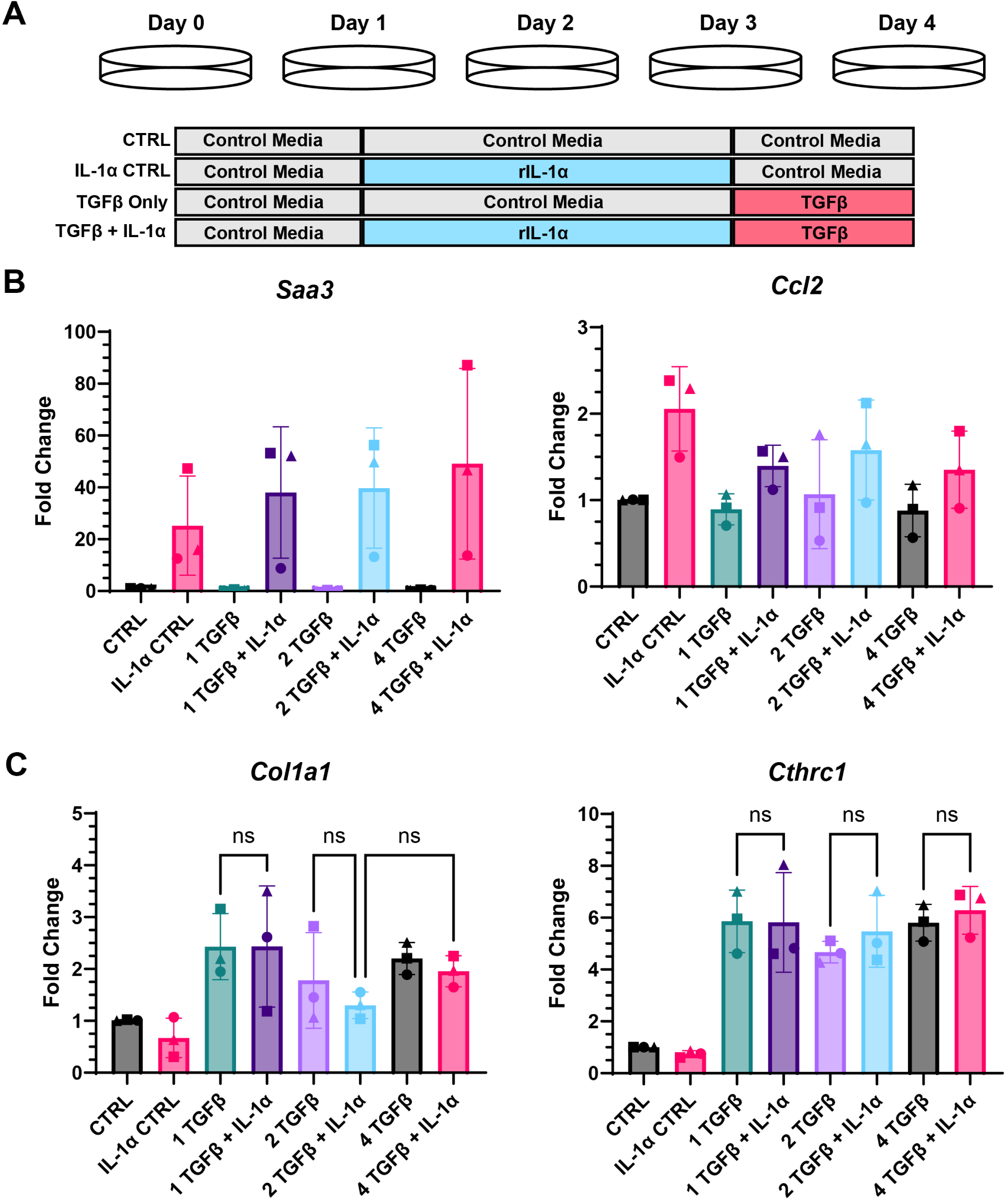
IL-1α-Driven Inflammatory Fibroblasts do not Exhibit Increased Sensitivity to TGFβ-Induced Fibrotic Stimulation. **(A)** Graphic summarizing the cell culture strategy employed to probe whether pre-treatment with 25pg/mL IL-1α primed fibroblasts to assume a fibrotic phenotype in response to TGFβ. Media was replaced every 24 hours. **(B)** *Saa3* and *Ccl2* gene expression by RT-qPCR for fibroblasts cultured on 0.5kPa plates under the conditions in (A). Media was supplemented with either 1, 2, or 4 ng/mL TGFβ. **(C)** *Col1a1* and *Cthrc1* gene expression by RT-qPCR for fibroblasts cultured on 0.5kPa plates under the conditions in (A). Media was supplemented with either 1, 2, or 4 ng/mL TGFβ. [(B) and (C)] Data shows mean fold change ± SD and are pooled from 3 independent experiments (n=3 mice; square, triangle, or circle). *P* values were calculated using a one-way ANOVA with post-hoc Turkey multiple comparison test: ns: not significant.

**Supplemental Figure 7:**
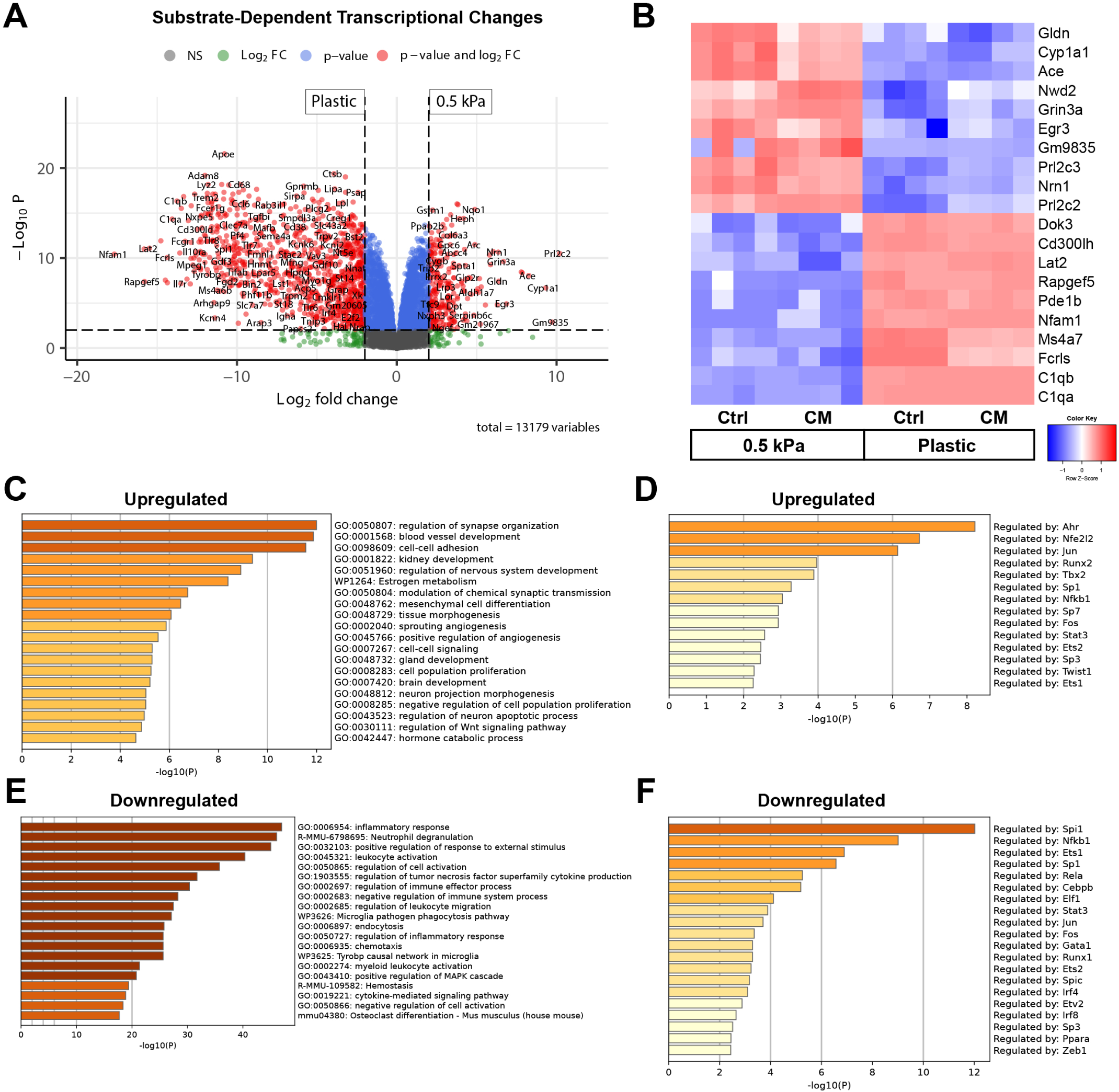
Culture Stiffness Induces Large-Scale Transcriptional Changes in Pulmonary Fibroblasts. **(A)** Volcano plot illustrating significantly differentially expressed genes between fibroblasts cultured on 0.5kPa or plastic plates treated with control and conditioned media with genes labeled. *P* value < 0.01, Fold Change > 272 upregulated 963 downregulated. **(B)** Heatmap depicting the relative expression of the top and bottom 10 most differentially expressed genes in (A). **(C)** Functional Enrichment analysis of the significantly upregulated genes in (A) via Metascape. **(D)** Enrichment analysis of transcription factors (TRRUST database via Metascape) using the significantly upregulated genes in (A). **(E)** Functional Enrichment analysis of the significantly downregulated genes in (A) via Metascape. **(F)** Enrichment analysis of transcription factors (TRRUST database via Metascape) using the significantly downregulated genes in (A).

